# Increased dosage of wild-type KRAS protein drives *KRAS*-mutant lung tumorigenesis and drug resistance

**DOI:** 10.1101/2024.02.27.582346

**Authors:** Tonci Ivanisevic, Wout Magits, Yan Ma, Peihua Zhao, Emiel Van Boxel, Benoit Lechat, Edward Stites, Greetje Vande Velde, Raj Sewduth, Anna Sablina

## Abstract

Almost 30% of lung adenocarcinomas are driven by activating *KRAS* mutations. The heterogeneous clinical behavior observed in these cancers could be due to the imbalance of wild-type and oncogenic KRAS alleles. However, the role of RAS dysregulation at the protein level needs to be further explored. A genome-wide global protein stability screen identified the CUL3 ubiquitin ligase adaptor LZTR1, as a major proteostatic regulator of wild-type but not mutant KRAS. In *KRAS*-mutant lung adenocarcinoma, shallow deletion of *LZTR1* is observed in up to 50% of patients and is associated with hypoxic signatures and poorer progression-free disease survival. In a *Kras*-mutant lung cancer mouse model, heterozygous loss of *Lztr1* promoted tumor growth, led to peritumoral vascular remodeling, and limited the response to the KRAS-G12D inhibitor MRTX1133. The vascular alteration in *LZTR1*-depleted lung cancer was mediated by increased wild-type KRAS protein dosage, which promoted mTOR pathway activation and a subsequent increase in VEGFA secretion. The inhibition of the PI3K/mTOR pathway using dactolisib normalized tumor vasculature, improved drug delivery, and overcame resistance to KRAS-G12D inhibitors. In summary, the dysregulation of RAS proteostasis contributes to lung tumorigenesis, and targeting wild-type KRAS signaling is crucial to overcome intrinsic resistance to inhibitors of mutant KRAS.

**Significance:** Our study highlights the impact of RAS proteostasis on lung cancer development and progression. Vascular normalization achieved by suppressing wild-type KRAS signaling improved the delivery and response to MRTX1133 in *KRAS*-mutant lung cancer with shallow *LZTR1* deletion.

## Introduction

Lung cancer is one of the most frequent cancers worldwide, with an aggressive clinical course and high mortality rates due to the lack of efficient long-term therapy. Almost 30% of non-small cell lung cancer (NSCLC) are driven by an activating Kirsten rat sarcoma viral oncogene homolog (*KRAS*) mutation ^1^. KRAS is a member of the RAS subfamily of small GTPases which, in addition to its founding members (KRAS4A, KRAS4B, NRAS, and HRAS), includes 35 RAS-related proteins ^2^. Mutations in the *KRAS* gene occur most frequently at codons G12, G13, and Q61, and activate KRAS signaling by either impairing the intrinsic GTPase activity of KRAS or structurally altering KRAS, to make it insensitive to GTPase-activating proteins ^3^.

In addition to oncogenic mutations, the dosage of mutant KRAS and wild-type RAS alleles due to copy number alterations (also called “RAS allelic imbalance”) also impacts RAS-mediated signaling and modulates drug sensitivity, leading to heterogeneous phenotypic characteristics and clinical outcomes. While the impact of a higher mutant *KRAS* allele dosage in lung cancer is well-defined, the significance of wild-type RAS allele dosage in tumorigenesis is still controversial. Both tumor-suppressive and tumor-promoting properties have been reported for the wild-type RAS proteins in the *KRAS*-mutant context ^4–8^. The increased dosage of wild-type RAS-like-without-CAAX-1 (RIT1) partially phenocopies KRAS in the lung cancer model ^9^, even though its activity depends on the classical RAS proteins ^10^. Mutant KRAS also leads to an increased expression of MRAS, which is part of a phosphatase complex that cooperates with RAS proteins for efficient MAPK pathway activation ^11^. Moreover, the activation of wild-type RAS signaling can drive therapeutic resistance to KRAS inhibitors in *RAS*-mutated cancers ^12^. However, the exact role of RAS protein dosage on tumorigenesis and drug response remains unexplored.

We and others have identified leucine zipper-like post-translational regulator 1 (LZTR1), a substrate adapter for the E3 ubiquitin ligase Cullin 3 (CUL3), as a proteostatic regulator of several RAS-like GTPases, including KRAS, HRAS, NRAS, MRAS, and RIT1 ^13–16^. It has been proposed that LZTR1-mediated ubiquitination of RAS could regulate its activity through either degradative or non-degradative mechanisms ^13,16^. In the wild-type RAS background, LZTR1-mediated RAS ubiquitination suppresses mitogen-activated protein kinase (MAPK) signaling and impairs cell growth ^13–16^.

Germline mutations concurrent with loss of heterozygosity at *LZTR1* are linked to several diseases, including glioblastoma and schwannomatosis ^17–19^. *LZTR1* mutations also predispose to pediatric neoplasms and are frequently observed in liver and testicular cancers. Allelic loss of *LZTR1* is also a common event in cancer. The Genomic Identification of Significant Targets in Cancer (GISTIC) analysis also demonstrated that focal deletions of *LZTR1* are observed in lung adenocarcinoma, pancreatic adenocarcinoma, and glioblastoma (https://portals.broadinstitute.org/tcga/home) ^20^. Furthermore, the *LZTR1* gene contains one of the most frequently retained introns across all cancers ^20,21^, further showing its potential role in tumorigenesis.

The full-body knock-out of *Lztr1* is embryonically lethal, whereas the heterozygous loss of *Lztr1* in mice partially recapitulates Noonan syndrome phenotypes, without leading to spontaneous tumorigenesis ^22^. Conditional inactivation of *Lztr1* and *Cdkn2a* in the mouse nervous system or *Lztr1* and *Lats1/2* in the Schwann cells promotes tumors in the peripheral nervous system, partially recapitulating schwannoma ^23,24^. Even though functional studies explicitly point out the tumor-suppressive role of *LZTR1* in wild-type RAS cancer, there is no evidence of the role of LZTR1 in the context of mutant KRAS. Here, we investigated how the dysregulation of RAS proteostasis and changes in RAS protein dosage contribute to the onset, progression, and treatment response of KRAS-mutant lung cancer.

## Results

### LZTR1 differentially regulates wild-type and mutant RAS proteins

Recent studies demonstrate that LZTR1 regulates the proteostasis of wild-type RAS-like GTPases ^14,16,25^, whereas its role in the regulation of KRAS oncogenic mutants is underexplored. To gain a better understanding of KRAS stability regulation, we performed a genome-wide global protein stability (GPS) screen ^26^, using a RAS stability reporter system, in which KRAS-WT or the KRAS-G12V mutant were fused to EGFP. The expression levels of the KRAS fusions relative to a dsRed control expressed from the same construct allowed us to infer the stability of the fused protein (Figure 1A).

**Figure 1.**
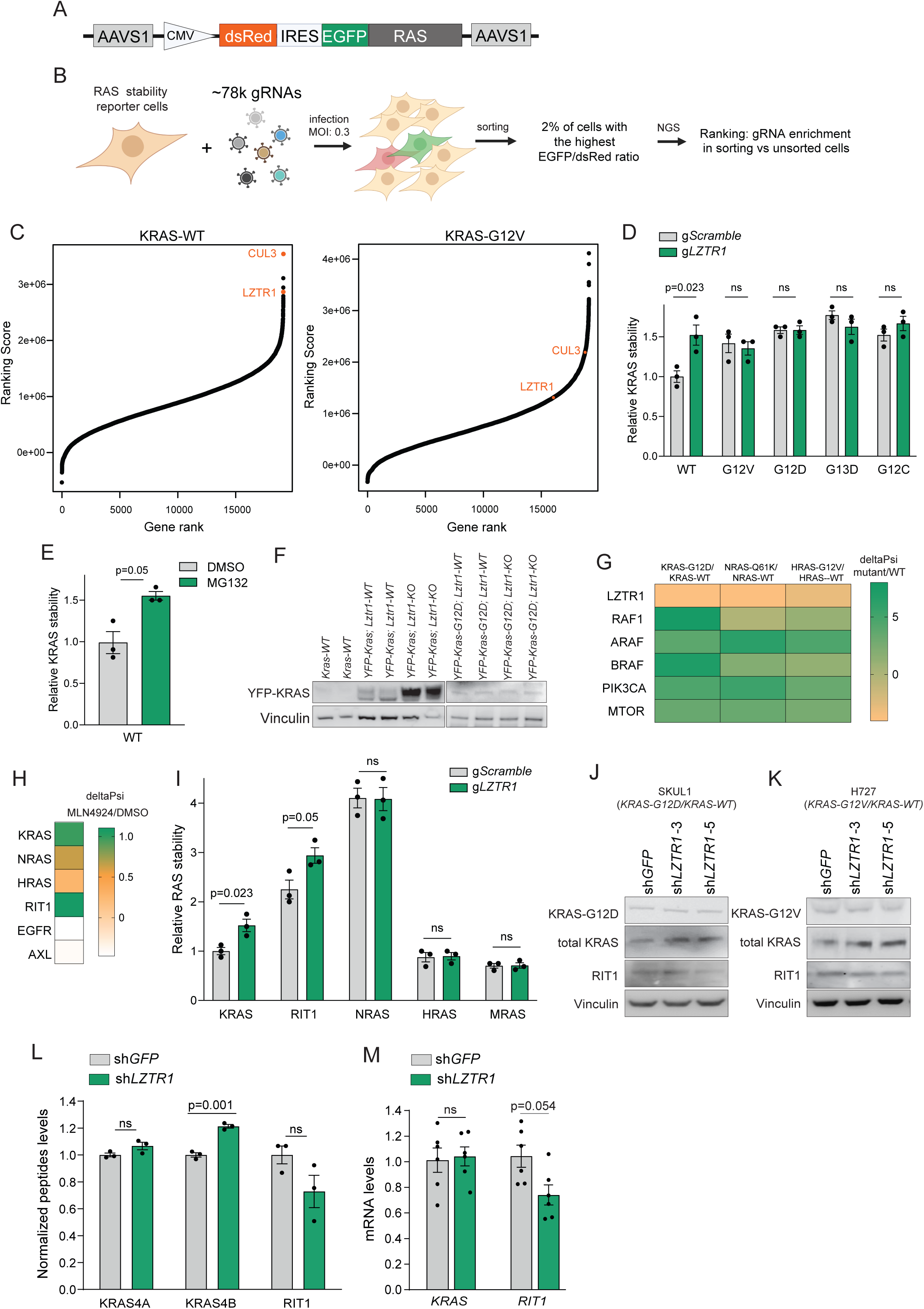
LZTR1 regulates the degradation of wild-type KRAS and RIT1. (A) The GPS cassette with KRAS-WT or KRAS-G12V was integrated into the AAVS1 locus of HAP1 cells. (B) Scheme representing the GPS-based CRISPR screen to identify regulators of KRAS-WT and KRAS-G12V stability. (C) The genome-wide GPS-based CRISPR screen identified regulators of KRAS-WT and KRAS-G12V stability. n=3. The ranking score of each of the genes was based on the gRNA enrichment score in sorted *vs* unsorted cells. (D) The GPS-based analysis of KRAS-WT and mutant KRAS protein stability in HEK293T cells expressing gScramble or gLZTR1. Data are shown as Mean±SEM, n=3. The p-value was calculated by a two-sided t-test. (E) The GPS-based analysis of KRAS-WT protein stability in HEK293T cells treated with DMSO or MG132 (10 μM, 24 hours). Data are shown as Mean±SEM, n=3. The p-value was calculated by a two-sided t-test. (F) Immunoblot analysis of YFP-KRAS-WT and YFP-KRAS-G12D in EPCAM-positive lung epithelial cells isolated from the indicated mice. (G) A heatmap showing the ratio of the BioIDBirA-fused RAS proteins. The BioID signals were obtained from ^51^. (H) The effect of MLN4924 treatment on the stability of wild-type RAS GTPases using GPS-based assays. A heatmap shows the ratios of dsRed/EGFP between DMSO or MLN4924 treated HEK293T cells. The ratios were obtained from ^29^. (I) The GPS-based analysis of the RAS protein stability in HEK293T cells expressing gScramble or gLZTR1. Data are shown as Mean±SEM, n=3. The p-value was calculated by a two-sided t-test. (J, K) Immunoblot analysis of KRAS and RIT1 expression in SKUL1 and H727 cells expressing shGFP or shLZTR1 using the indicated antibodies. (L) TMT-labeled proteomic analysis of H727 cells expressing shGFP or shLZTR1. Quantification of peptides specific to KRAS and RIT1 is shown as Mean±SEM, n=3. The p-value was calculated using a two-sided t-test. (M) RT-qPCR analysis of *KRAS* and *RIT1* expression in H727 cells expressing shGFP or shLZTR1. Data are shown as Mean±SEM, n=6. The p-value was calculated by a two-sided t-test.

To obtain the physiological level of tagged KRAS, the GPS constructs were integrated into the AAVS1 locus of HAP1 cells using a CRISPR-based approach (Figure 1A). We next selected single-cell clones, which showed near-endogenous EGFP-KRAS or EGFP-KRAS-G12V levels, and then overexpressed CAS9 in the selected clones (Figure S1A, B). The generated GPS reporter cells were then infected with the Brunello genome-wide CRISPR knockout library with an MOI of 0.3. 5 days after the introduction of the gRNA library, we sorted the cells with the top 2% highest EGFP/dsRed ratio and purified the DNA from unsorted and sorted populations followed by PCR amplification of the gRNA sequence and next-generation sequencing (Figure 1B). We ranked the genes according to the gRNA enrichment or abundance in the sorted *vs* unsorted populations. Both LZTR1 and CUL3 were the top hits of the GPS screen for KRAS-WT (Figure 1C). Four different gRNAs targeting *LZTR1* showed consistent enrichment in the sorted fraction with a high EGFP to dsRed ratio (Figure S1C). No other E3 ubiquitin ligase scored high in the GPS screen, implicating the LZTR1/CUL3 complex as the main regulator of KRAS-WT stability. On the other hand, neither LZTR1 nor CUL3 scored in the GPS screen for KRAS-G12V (Figure 1C), suggesting that oncogenic KRAS mutations abrogate LZTR1/CUL3-mediated degradation.

We confirmed the results of the GPS screen in a low-throughput format. The loss of LZTR1 increased the stability of KRAS-WT, to a level comparable to the treatment with the proteasomal inhibitor MG132 (Figure 1D, E), indicating a key role of LZTR1 in controlling the proteasomal degradation of KRAS-WT. Conversely, all tested oncogenic KRAS mutants, G12V, G12C, G12D, and G13D, exhibited higher stability when compared to KRAS-WT and were not stabilized by *LZTR1* knockdown (Figure 1D). We also assessed the effect of *Lztr1* loss in a reporter mouse model expressing either KRAS-WT or KRAS-G12D with an endogenously inserted YFP tag ^27^. We observed increased protein expression of KRAS-WT in isolated lung epithelial cells from *Lztr1*-KO mice, whereas *Lztr1* deletion did not affect the expression of the YFP-KRAS-G12D protein (Figure 1F). Concordantly, the DepMap analysis of lung cancer cell lines revealed that the expression of LZTR1 showed a strong negative correlation with the expression levels of KRAS protein in lung cancer cell lines with KRAS-WT, whereas cell lines harboring mutant *KRAS* did not show a strong correlation (Figure S1D). Furthermore, a previous study ^28^ showed that, unlike downstream effectors, LZTR1 showed a higher affinity to RAS-WT proteins when compared to oncogenic RAS mutants (Figure 1G). Altogether these results indicate that, whereas LZTR1 controls the expression levels of KRAS-WT, it does not affect the stability of oncogenic *KRAS* mutants.

A recent genome-wide study ^29^ revealed that RAS-like GTPases showed different levels of stabilization in response to the MLN4924 inhibitor, which blocks the activity of the Cullin-RING ubiquitin ligases (CRL) (Figure 1H). At the same time, MLN4924 treatment did not affect the stability of EGFR or AXL, which were also proposed as LZTR1 substrates ^24^. This suggests that LZTR1 differentially affects the degradation of RAS-related GTPases while questioning the contribution of the LZTR1/CUL3 in the regulation of the EGFR or AXL stability. Similar to CRL inhibition ^29^, *LZTR1* knockout by gRNA led to increased stability of KRAS and RIT1 but did not affect NRAS, HRAS, or MRAS stability (Figure 1I).

These results suggest that LZTR1 loss would lead to increased expression of wild-type KRAS and RIT1. Immunoblot analysis using antibody specific to mutant KRAS showed similar levels of mutant KRAS protein in H727 (*KRAS-G12V* and *KRAS-WT* alleles) and SKUL1 (*KRAS-G12D* and *KRAS* alleles-WT) lung cancer cells expressing either sh*GFP* or sh*LZTR1*. On the other hand, total levels of KRAS (mutant and WT) were increased in both cell lines, as detected by immunoblotting with a pan-KRAS antibody (Figure 1J, K, Figure S1E). TMT-labeled proteomics confirmed that total KRAS (G12V and WT) levels were significantly higher upon *LZTR1* loss in lung cancer H727 cells (Figure 1L). The elevated protein expression of KRAS could be attributed to increased protein stability, as we did not observe any changes in *KRAS* mRNA levels (Figure 1M). Even though LZTR1 loss leads to RIT1 stabilization, we did not observe an increase of RIT1 protein expression in LZTR1-depleted lung cancer cells which could be due to the decreased expression of *RIT1* at mRNA level. In conclusion, LZTR1 differentially affects the dosage of wild-type RAS-related GTPases in the *KRAS*-mutant context.

### LZTR1 haploinsufficiency promotes KRAS-driven lung cancer and resistance to targeted therapy

We next investigated the role of LZTR1-mediated regulation of RAS proteostasis in KRAS-driven tumorigenesis. The Cancer Genome Atlas (TCGA) analysis of the lung adenocarcinoma (LUAD) dataset showed that heterozygous loss of *LZTR1*-containing locus led to decreased *LZTR1* expression and was observed in about 50% of *KRAS*-mutant lung adenocarcinoma (Figure 2A, B). Heterozygous loss of *LZTR1* or mutations significantly co-occurred with oncogenic *KRAS* mutations and were associated with poorer progression-free disease survival (Figure 2A, C), suggesting that *LZTR1* loss contributes to tumorigenesis driven by mutant KRAS.

**Figure 2.**
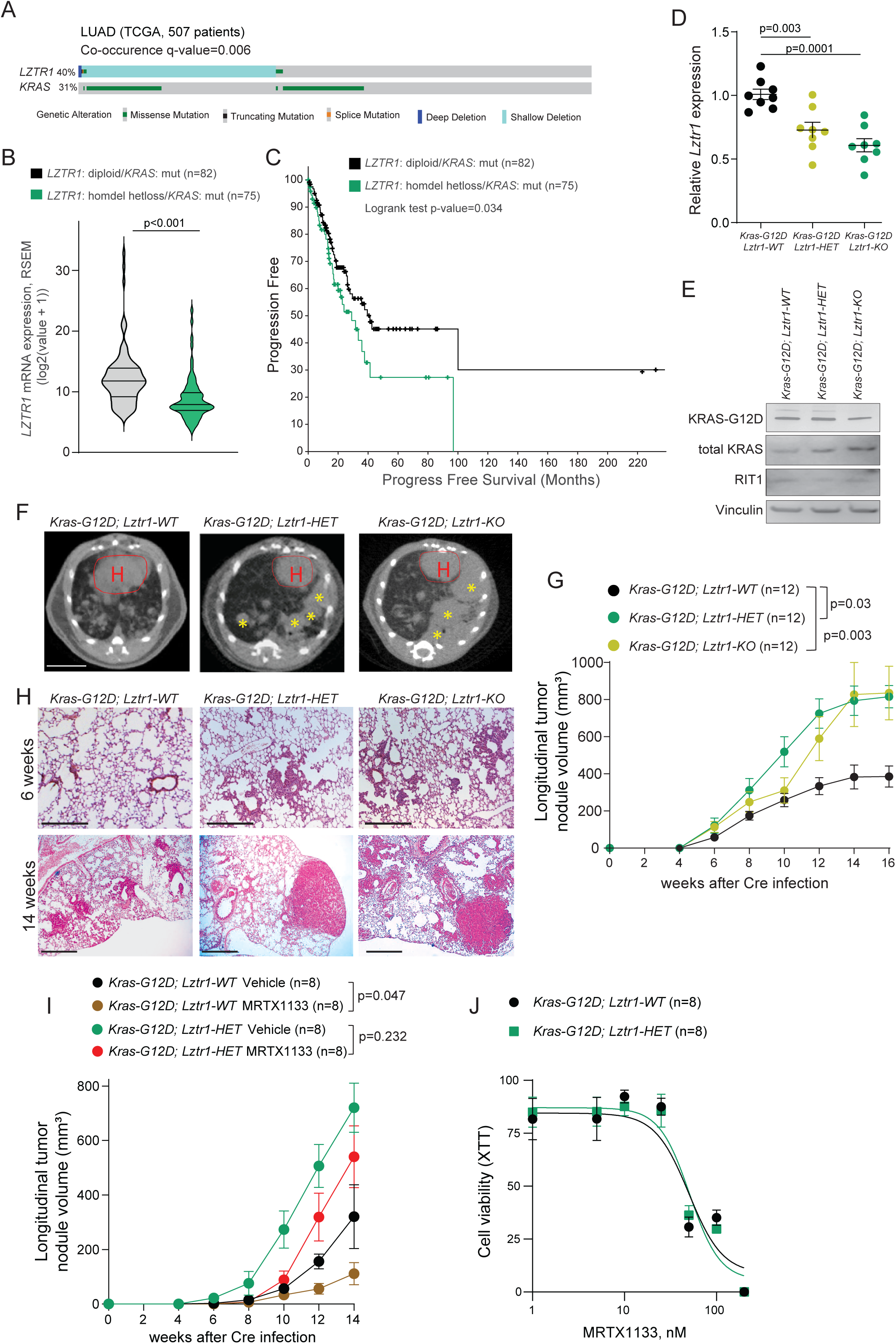
*Lztr1* loss facilitates Kras G12D-driven lung tumorigenesis. (A) The co-occurrence of *KRAS* mutation and *LZTR1* alterations in TCGA lung adenocarcinoma (LUAD) PanCancer dataset (n=507 patients) obtained from cBioPortal. (B) *LZTR1* expression in TCGA LUAD tumors with different *LZTR1* statuses. Homdel, homozygous deletion; hetloss, shallow deletion. The p-value was calculated by cBioPortal. (C) Progression-free survival of KRAS-mutant TCGA LUAD patients with different *LZTR1* statuses. Homdel, homozygous deletion; hetloss, shallow deletion. The p-value was calculated by cBioPortal. (D) RT-qPCR analysis of *Lztr1* expression in lung tumor cells isolated from mice with the indicated genotype 14 weeks after injection with adenovirus particles expressing *Sftpc-*specific Cre. Data are shown as Mean±SEM, n=8 per group, p-value was calculated by the Mann-Whitney test. (E) Immunoblot analysis of KRAS and RIT1 expression in lung tumor cells isolated from ice with the indicated genotype 14 weeks after injection with adenovirus particles expressing *Sftpc-*specific Cre recombinase. (F) Representative reconstructed micro-Computed Tomography (micro-CT) scans of the lungs of mice with the indicated genotype 14 weeks after Cre injection. The yellow asterisk indicates tumor nodules, “H” and the red line delineate the heart. Scale bar: 1 cm (G) Longitudinal quantification of the tumor nodule growth assessed by tumor volumetric analysis of the micro-CT scans obtained at different timepoints after Cre injection. Tumor volumetric analysis was performed by segmentation and feature extraction using MIPAV (NIH). Data are shown as Mean±SEM; n=12 per group. The p-value was calculated by Mixed model analysis. (H) Hematoxylin-Eosin (H&E) staining of the lung tissues of the mice with the indicated genotypes 6 and 14 weeks after Cre injection. Scale bar, 100 µm. (I) Quantification of total tumor nodule volume using micro-CT scans of the lung of mice treated with vehicle or MRTX1133 (10 mg/kg, intraperitoneal injection every 2 days) after tumor nodule delineation. Data are shown as MEAN±SEM; n=8 per group. The p-value was calculated using Mixed model analysis. (J) The growth of lung cancer cells isolated from mice with the indicated genotypes treated with increasing doses of MRTX1133 measured using XTT assay. Data are shown as MEAN±SEM; n=8 per group.

To assess the impact of *Lztr1* loss on *KRAS*-driven lung cancer in a system that recapitulates LUAD ^30^, we used the *Kras^G12D^ ^lsl/wt^* mouse model that developed lung adenocarcinoma after intratracheal injection of adenovirus expressing Cre-recombinase driven by the surfactant protein C (*Sftpc*) promoter, specific to the alveolar type II epithelial cells. We confirmed suppression of *Lztr1* expression and induction of KRAS-G12D expression in sorted lung tumor cells isolated after Cre injection (Figure 2D, E). Consistently to human lung cancer cells, immunoblot analysis revealed no differences in KRAS-G12D levels or RIT1 in cells with different *Lztr1* status, whereas total KRAS levels (wt and KRAS-G12D) were increased in cells with *Lztr1* loss (Figure 2E).

To assess the role of *Lztr1* deletion in *Kras*-mutant lung cancer, we monitored the growth of *Lztr1^wt/wt^, Kras^G12D^ ^lsl/wt^* (*Kras-G12D*; *Lztr1-WT*); *Lztr1^flox/wt^, Kras^G12D^ ^lsl/wt^* (*Kras-G12D*; *Lztr1-HET*) and *Lztr1^flox/flox^, Kras^G12D^ ^lsl/wt^* (*Kras-G12D; Lztr1-KO)* tumors, using micro-computed tomography (micro-CT) (Figure 2F). The volumetric tumor analysis demonstrated that either *Lztr1-KO* or *Lztr1-HET* mice showed an increased tumor burden (Figure 2G). Moreover, hematoxylin-eosin (H&E) staining revealed that the lesions appeared in the *Kras-G12D; Lztr1-HET or Kras-G12D; Lztr1-KO* lungs at an earlier stage, progressing rapidly into large, compacted adenoma nodules (Figure 2H). Further immunohistochemistry analysis of the lung tissue revealed that the tumor lesions showed positivity for GATA6, cytokeratin 19 (CK19), and transcription termination factor 1 (TTF1) (Figure S2), indicating that the mouse model recapitulates characteristics of human papillary and glandular lung adenocarcinomas.

Thus, loss of either one or two *Lztr1* alleles induced earlier tumor onset and accelerated tumor progression, indicating that *LZTR1* acts as a haploinsufficient tumor suppressor in the context of *KRAS*-mutant lung cancer. These results also suggest an oncogenic role of increased wild-type KRAS dosage. In line with this idea, a mathematical model of RAS signaling studying the interplay between mutant RAS and wild-type RAS-GTP ^31–33^ revealed that a 50% increase in wild-type KRAS results in an 8% increase in total GTP-bound RAS-proteins, suggesting that the changes in wild-type KRAS abundance could further potentiate oncogenic signaling.

Increased activity of wild-type RAS proteins has been also reported to result in acquired resistance to the inhibitors targeting mutant KRAS ^34–36^. Therefore, we assessed how *Lztr1* loss can affect response to MRTX1133, a small-molecule, non-covalent, and selective KRAS-G12D inhibitor. MRTX1133 has demonstrated potent *in vitro* and *in vivo* antitumor efficacy against KRAS-G12D-mutant cancer cells ^37^. We found that whereas MRTX1133 suppressed the growth of *Kras-G12D; Lztr1-WT* tumors, it only moderately delayed tumor growth of *Kras-G12D; Lztr1-HET* tumors at earlier stages and did not show a significant effect at later timepoints (Figure 2I). This indicates that the heterozygous loss of *LZTR1* commonly observed in LUAD patients could explain intrinsic resistance to the inhibitors targeting mutant KRAS.

Furthermore, *in vitro*, isolated *Lztr1-WT* and *Lztr1-HET* tumor cells showed similar sensitivity to the MRTX1133 treatment (Figure 2J), suggesting that alterations in the tumor microenvironment could be responsible for the limited response of *Lztr1-HET* tumors to KRAS-G12D inhibition.

### LZTR1 haploinsufficiency leads to remodeling of tumor-associated vasculature

The gene set enrichment analysis (GSEA) of *KRAS*-mutant LUADs with different *LZTR1* copy number variation status revealed that *LZTR1* loss was associated with hypoxic gene expression signature (Figure 3A). *Lztr1* haploinsufficiency in the *Kras*-mutant mouse lung cancer model led to increased hypoxia as shown by increased HIF1α expression (Figure 3B). Therefore, we analyzed the tumor-adjacent vasculature in the lungs of *Kras-G12D; Lztr1-WT* and *Kras-G12D; Lztr1*-HET mice. Optical tomography of cleared lungs revealed a significant difference between the blood vessels surrounding the tumors in *Kras-G12D; Lztr1-HET* and *Kras-G12D; Lztr1-WT* mice (Figure 3C). In *Lztr1-WT* mice, the blood vessels had a clear and organized structure, with larger vessels that branched into smaller ones. The tumor associated vasculature in the *Lztr1-HET* mice displayed disorganization and a lack of conventional hierarchy of blood vessels, typical for tumor vasculature ^38^. The tumor-adjacent vessels of the *Lztr1-HET* mice were also leaky, according to FITC-dextran diffusion assay (Figure 3D). In *Lztr1-HET* mice, pronounced ICAM1 expression in endothelial cells and a higher count of CD11-positive immune cells at vascular junctions also indicated vascular inflammation (Figure 3E). Moreover, SMA immunostaining revealed reduced pericyte coverage of the *Lztr1*-*HET* vasculature, explaining the increased vascular permeability and remodeling (Figure 3F). Altogether, these results show that *Lztr1* haploinsufficiency in cancer cells drives aberrant vasculature in the tumor and surrounding tissue.

**Figure 3.**
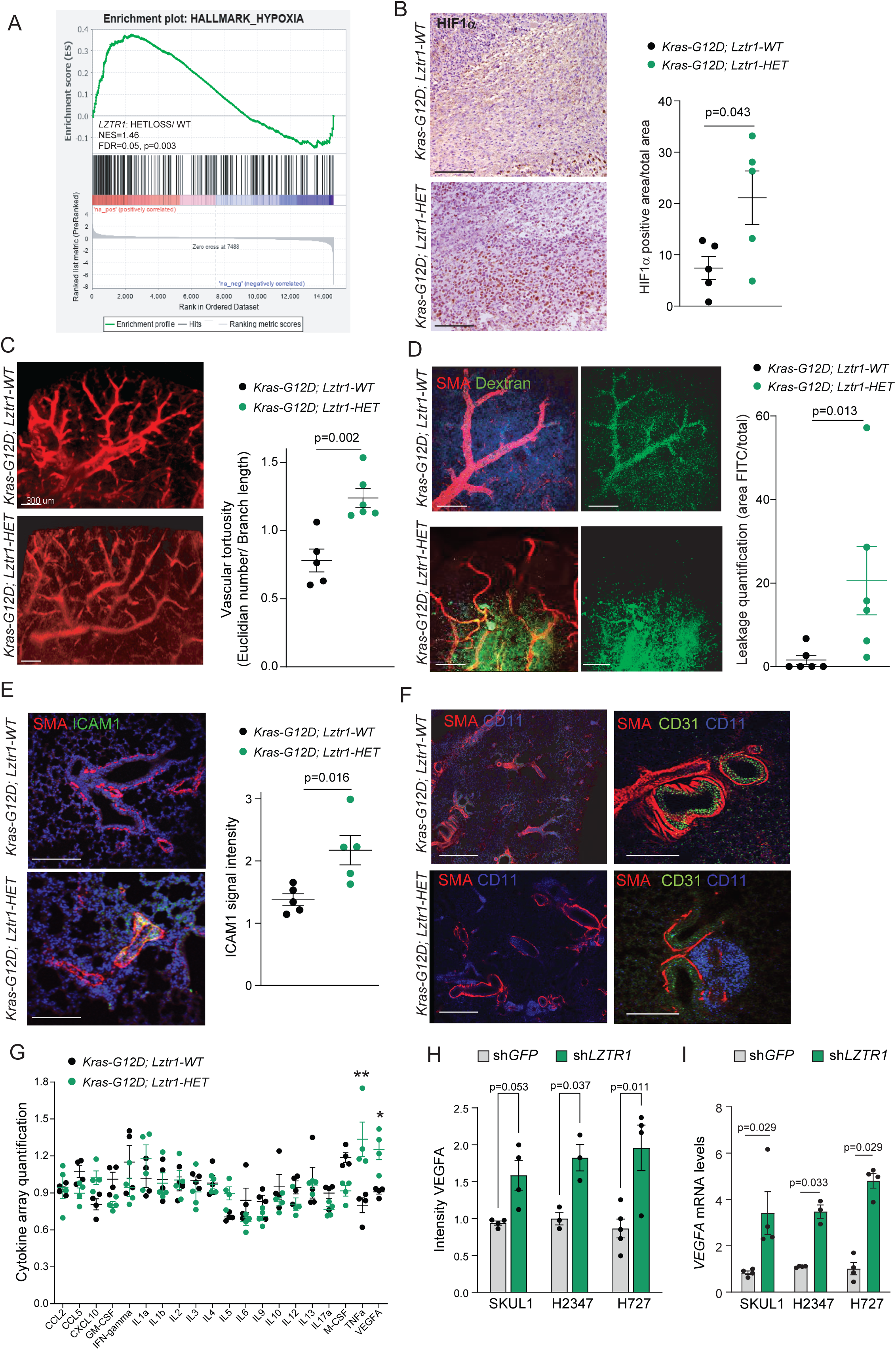
Lztr1 loss in *Kras*-mutant tumors leads to remodeling of the tumor vasculature. (A) Gene Set Enrichment Analysis (GSEA) of the hypoxia pathway signature in LUAD patients with different *LZTR1 status*. (B) Immunostaining analysis of HIF1α in lung adenoma nodules in mice with the indicated genotypes, at 14 weeks after Cre injection. Scale bar: 30 µm. Quantification of HIF1α-positive to total area using ZEN Axiovision. Data are shown as Mean±SEM; n=5 per group. The p-value was calculated by the Mann-Whitney test. (C) Optical projection tomography of the tumor-adjacent arterial network in the indicated mice after SMA immunostaining. Scale bar, 300 µm. Quantification of vascular tortuosity after reconstruction of the vascular network was performed as described in Material and Methods. Data are shown as Mean±SEM; n=5 per group. The p-value was calculated by the Mann-Whitney test. (D) Imaging of the tumor-adjacent vascular leakage after FITC-Dextran injection and SMA immunostaining in mice with the indicated genotype. Scale bar, 50µm. Quantification of vascular leakage is shown as Mean±SEM; n=5 per group. The p-value was calculated by the Mann-Whitney test. (E) Immunostaining analysis of the tumor-adjacent vasculature of mice with the indicated genotype using anti-ICAM1 and anti-SMA antibodies. Scale bar, 50µm. Quantification of ICAM-positive is shown as Mean±SEM; n=5 per group. The p-value was calculated by the Mann-Whitney test. (F) Immunostaining analysis of the tumor-adjacent vasculature of mice with the indicated genotype using CD11, SMA, and CD31 specific antibodies. Scale bar, 100µm. (G) Cytokine array assay of the supernatant of lung cancer cells isolated from the indicated mice. n=4 per group. Data are shown as Mean±SEM; n=4 per group. The p-value was calculated by the Mann-Whitney test. (H) ELISA-based quantification of VEGFA levels in the supernatant of *KRAS*-mutant human lung cancer cell lines. Data are shown as Mean±SEM; n=4 per group. The p-value was calculated by the Mann-Whitney test. (I) RT-qPCR analysis of VEGFA expression in *KRAS*-mutant human lung cancer cell lines. Data are shown as Mean±SEM; n=4 per group. The p-value was calculated by the Mann-Whitney test.

To get a better understanding of the paracrine effect of *Lztr1* loss, we performed a secretome analysis using the supernatants of lung cancer cells isolated from *Lztr1-WT* and *Lztr1*-*HET* mice. The ELISA-based cytokine array analysis revealed that vascular endothelial growth factor A (VEGFA) and tumor necrosis factor α (TNFα) were significantly upregulated in the supernatant of *Lztr1-HET* cells (Figure 3G). Consistently, *LZTR1* depletion in human lung cancer cells also resulted in increased VEGFA secretion (Figure 3H). Increased VEGFA secretion by *LZTR1*-depleted cells could be explained by increased *VEGFA* mRNA levels (Figure 3I). These results suggest that increased VEGFA expression by tumors with *LZTR1* loss might lead to vasculature abnormalities.

### Increased wild-type KRAS protein dosage leads to aberrant vasculature

The gene set enrichment analysis (GSEA) of *KRAS*-mutant TCGA LUADs with or without *LZTR1* shallow deletion revealed that *LZTR1* loss in lung adenocarcinoma patients was associated with mammalian target of rapamycin (mTOR) pathway-related gene expression signatures (Figure 4A). Concordantly, the integrated results of proteome and phosphoproteome analysis of human lung cancer cells expressing sh*GFP* or sh*LZTR1* demonstrated that mTOR signaling was one of the top-altered signaling pathways (Figure 4B, C). Immunoblotting analysis confirmed that suppression of *LZTR1* in human lung cancer cells, SKUL1 and H727, led to increased phosphorylation of the mTOR branch of the RAS pathway (Figure 4D).

**Figure 4.**
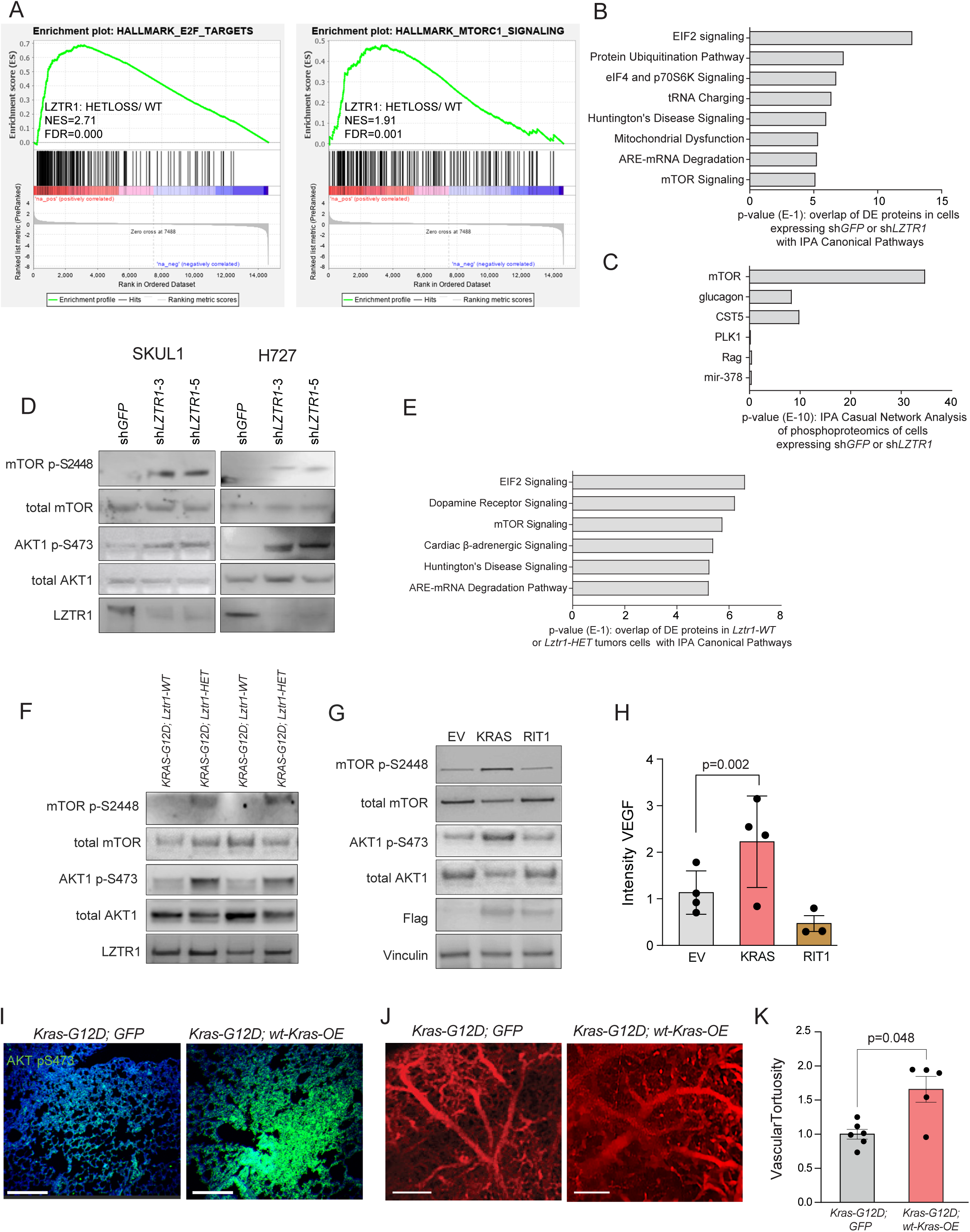
The impact of Lztr1 deletion on RAS signaling. (A) Gene Set Enrichment Analysis of the mTOR pathway-related signatures in LUAD patients with different *LZTR1* status. (B) The Canonical Pathway analysis of Ingenuity Pathway Analysis (IPA, QIAGEN) of differentially expressed (DE) proteins in H727 cells expressing sh*GFP* or sh*LZTR1* identified by TMT-labeled MS proteomics, n=3 per group. (C) The Causal Network IPA analysis of differentially phosphorylated proteins of H727 cells expressing sh*GFP* or sh*LZTR1* identified by Ti-IMAC-enriched phosphoproteomics, n=3 per group. (D) Immunoblot analysis of phosphorylated and total AKT1 and mTOR in human lung cancer cells, SKUL1 and H727, expressing sh*GFP* or shRNAs against *LZTR1*. (E) The Canonical Pathway IPA analysis of differentially expressed (DE) proteins in lung cancer cells isolated from the indicated mice identified by TMT-labeled MS proteomics, n=3 per group. (F) Immunoblot analysis of phosphorylated and total AKT1 and mTOR in lung cancer cells isolated from *Kras-G12D; Lztr1-WT* and *Kras-G12D; Lztr1-HET* mice. (D) Immunoblot analysis of phosphorylated and total AKT1 and mTOR levels in SKUL1 cells overexpressing an empty vector (EV), KRAS-WT, or RIT1-WT. (H) ELISA-based quantification of VEGFA levels in the supernatant of SKUL1 cells expressing the indicated constructs. Data are shown as Mean±SEM; n=4 per group. The p-value was calculated by the Mann-Whitney test. (I) Immunostaining analysis of phosphorylated AKT1 and SMA on lung tumor sections of *Kras-G12D; Lztr1-WT* mice, 10 weeks after injection of a lentivirus expressing with Cre only or HA-KRAS and Cre under the control of the *Sftpc* promoter. (J) Optical projection tomography imaging of the tumor-adjacent arterial network of lungs of mice with *Kras-G12D; Lztr1-WT* 10 weeks after injection with lentivirus expressing Cre only or HA-KRAS and Cre under the control of the *Sftpc* Cre-promoter, 20 to 30% lethality observed in the HA-KRAS lentivirus injected group. The lungs were immunostaining with SMA-specific antibody. Scale bar, 300 µm. (J) Quantification of vascular tortuosity after reconstruction of the vascular network was performed as described in Material and Methods. Data are shown as Mean±SEM; n=5 per group. The p-value was calculated by the Kolmogorov-Smirnov test.

The proteome analysis of lung tumor cells isolated from *Lztr1-WT* and *Lztr1*-*HET* mice also revealed the alteration of mTOR signaling (Figure 4E). Immunohistochemistry and immunoblotting analysis of *Kras-*mutant isolated tumor cells also showed increased phosphorylation of AKT1 and mTOR, indicating upregulation of PI3K/mTOR signaling in the *Lztr1-HET* mice (Figure 4F, Figure S3A, B). In contrast, we detected only a minor increase of MEK1 or ERK1/2 phosphorylation in the isolated cells (Figure S3A, B). Given that *LZTR1* loss was previously shown to significantly increase the MAPK activity in RAS-WT cells ^13–16,39^, these results indicate that the LZTR1 effect on RAS signaling could be context-dependent.

As a next step, we evaluated whether activation of mTOR signaling could be attributed to LZTR1-mediated increased stability of KRAS and RIT1. For this, we overexpressed either KRAS-WT or RIT1 in SKUL1 cells harboring heterozygous *KRAS-G12D* mutation. Immunoblot analysis revealed that the overexpression of KRAS-WT, but not RIT1, led to increased phosphorylation of mTOR and AKT1 (Figure 4G). Moreover, KRAS-WT overexpression led to increased VEGFA secretion (Figure 4H). Furthermore, the injection of a lentivirus expressing both Cre and KRAS-WT under *Sftpc* specific promoter into *Kras^G12D^ ^lsl/wt^* (*Kras-G12D; WT-KRAS-OE)* mice led to increased activity of mTOR and vasculature remodeling (Figure 4I-K). Altogether, these results demonstrate that increased KRAS-WT expression triggered by loss of LZTR1 induces the activity of the PI3K/mTOR branch of the RAS signaling, leading to an increase of VEGFA secretion, and vascular alterations.

### Identification of therapies for normalization of vasculature

To discover therapies that could normalize vascular dysfunction driven by *LZTR1* loss, we screened for the RAS pathway inhibitors that could block VEGFA secretion in *LZTR1*-depleted lung cancer cells (Figure 5A). In line with the results of the signaling analysis, we found that the AKT inhibitor Ipasertib, the mTOR inhibitor Everolimus, and the dual PI3K/mTOR inhibitor dactolisib (NVP BEZ-235) rescued the VEGFA upregulation induced by *LZTR1* depletion (Figure 5A). These results indicate that increased secretion of VEGFA in lung cancer cells with *LZTR1* loss could be due to upregulation of the PI3K/mTOR pathway. On the other side, treatment with the KRAS-G12D inhibitor, MRTX1133 or MEK1/2 inhibitors, Selumetinib, and Trametinib, did not suppress enhanced VEGFA secretion triggered by *LZTR1* loss (Figure 5A).

**Figure 5.**
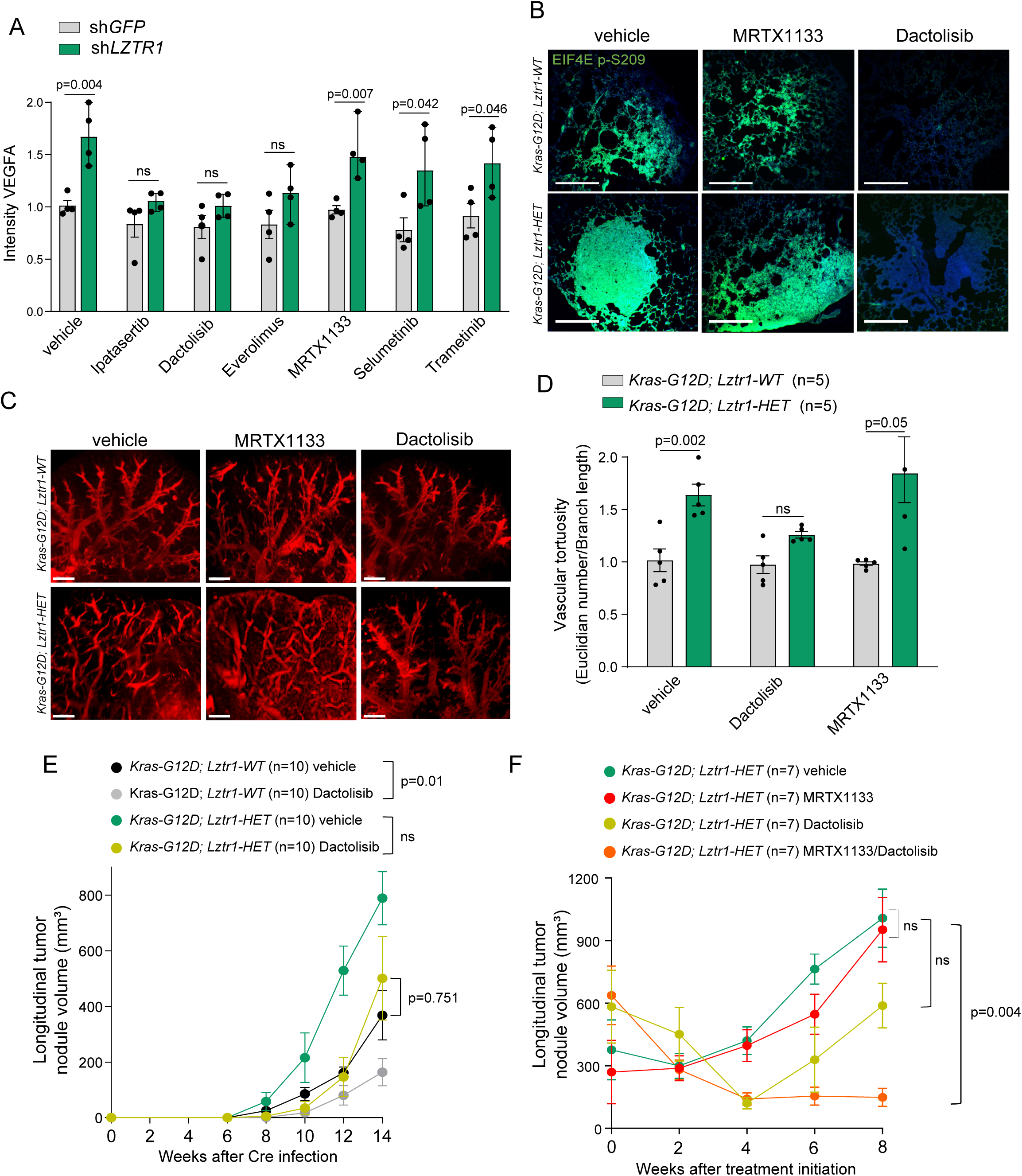
The combinatory treatment of MRTX1133 and dactolisib suppresses the growth of *Kras*-mutant tumors with Lztr1 loss. (A) ELISA-based quantification of VEGFA levels in the supernatant of H727 cells expressing shGFP or sh*LZTR1,* after treatment with the indicated drugs for 24 hours. Data are shown as Mean±SEM; n=4 per group. The p-value was calculated by the Mann-Whitney test. (B) Immunostaining analysis of phosphorylated EIF4E on lung tumor sections of the mice with the indicated genotypes, 14 weeks after the *Sftpc*-specific Cre injection and treatment with vehicle, dactolisib (10 mg/kg, IP every 2 days), or MRTX1133 (10 mg/kg, IP injection every 2days). Scale bar, 100µm. (C) Optical projection tomography imaging of the tumor-adjacent arterial network of mice with the indicated genotypes, 14 weeks after the *Sftpc*-specific Cre injection and treatment with dactolisib (10 mg/kg, IP every 2 days) or MRTX1133 (10 mg/kg, IP injection every 2days). The lungs were immunostaining with SMA-specific antibody. Scale bar, 300 µm. (D) Quantification of vascular tortuosity after reconstruction of the vascular network in the indicated mice. Data are shown as Mean±SEM; n=5 per group. The p-value was calculated by the Mann-Whitney test. (E) Quantification of total tumor nodule volume using micro-CT scan in the indicated mice treated with vehicle or dactolisib (10 mg/kg, IP injection every 2 days) after tumor nodule delineation. Data are shown as MEAN±SEM; n=10 per group. The p-value was calculated by Mixed model analysis. (F) Quantification of total tumor nodule volume using micro-CT scan in the indicated mice treated with vehicle, dactolisib (10 mg/kg, IP every 2 days), and/or MRTX1133 (10 mg/kg, IP every 2 days) after tumor nodule delineation, starting from week 12 after the *Sftpc-*specific Cre injection. Data are shown as MEAN±SEM; n=7 per group. The p-value was calculated by Mixed model analysis.

The administration of MRTX1133 also failed to suppress increased EIF4E phosphorylation in *Lztr1-HET* mice and did not lead to the normalization of the tumor-associated vasculature (Figure 5B-D). Conversely, inhibition of mTOR activity by dactolisib normalized the vasculature in *Lztr1-HET* mice (Figure B-D). These results further support the idea that the dysregulation of wild-type RAS protein dosage contributes to the upregulation of the mTOR pathway and abnormal vasculature in the LZTR1-depleted context. Moreover, dactolisib abolished the difference between the growth of *Lztr1-WT* and *Lztr1-HET* tumors (Figure 5E), suggesting that the loss of *Lztr1* facilitates tumorigenesis due to vasculature dysfunction caused by upregulation of the mTOR signaling pathways.

Given that *in vitro*, *Lztr1-WT* and *Lztr1-HET* tumor cells showed similar sensitivity to the MRTX1133 treatment (Figure 1I, J), this suggests that vascular normalization by dactolisib can overcome resistance to MRTX1133 treatment observed in *Lztr1-HET* mice. In agreement with this idea, we found that the combination of MRTX1133 and dactolisib showed promising long-term effects on the reduction of tumor growth in the preclinical model of lung cancer (Figure 5F). These results suggest that targeting of wild-type RAS signaling in lung cancer cells can enhance the efficacy of MRTX1133, a selective inhibitor of KRAS-G12D, by normalizing tumor-adjacent environment and improving drug delivery.

## Discussion

The dosage of either mutant or wild-type KRAS allele leads to high heterogeneity of *KRAS-*mutant lung tumors and challenges the attempts to rationally design drug combinations for lung cancer patients ^40,41^. In our study, we investigated the mechanisms governing the proteostatic regulation of mutant and wild-type RAS proteins. We also uncovered the impact of RAS protein stability on the growth and progression of *KRAS*-mutant lung cancer and the response to inhibitors targeting mutant KRAS.

A genome-wide GPS screen identified LZTR1 as a key regulator of wild-type KRAS stability, confirming previous observations that the LZTR1/CUL3 complex controls KRAS ubiquitination ^13–16^. The most common cancer-associated mutations of *KRAS* abrogate the ability of LZTR1 to regulate their degradation, leading to increased half-life of KRAS mutants when compared to their wild-type KRAS protein. Given that an increased dosage of mutant KRAS promotes tumorigenesis, this finding suggests an additional mechanism by which KRAS mutations contribute to its oncogenic properties. These results also indicate that *LZTR1* loss could contribute to the disease by promoting wild-type RAS signaling.

Several studies have previously demonstrated that LZTR1 regulates the ubiquitination of different RAS family members ^13–16,39^. However, the quantitative monitoring of RAS stability using the GPS approach revealed that LZTR1-mediated ubiquitination significantly affects only the stability of KRAS and RIT1. This suggests that while LZTR1 modulates KRAS and RIT1 by promoting proteasomal degradation, it could regulate the activity of other RAS family members through non-degradative mechanisms. Specifically, we previously found that LZTR1-mediated non-degradative ubiquitination of HRAS at K170 inhibits its activity by detaching the protein from the plasma membrane ^16^. On the other hand, the ubiquitination of the C-terminus hypervariable region of the KRAS ortholog leads to its degradation ^42^. This suggests that the differences in hypervariable regions might account for the different outcomes of LZTR1-mediated ubiquitination.

In our study, we identified *LZTR1* as a haploinsufficient tumor suppressor in the context of *KRAS*-mutant lung cancer. In mice, *Lztr1* haploinsufficiency also mimics some of the features of Noonan syndrome, such as craniofacial defects and cardiac abnormalities ^16^. Recent studies demonstrated that another CUL3 substrate adaptor SPOP forms high-order oligomers when its protein concentration reaches a certain threshold, a process crucial for its activity ^43^. SPOP oligomerizes due to the presence of the BTB-BACK domains, which resemble the BTB-BACK domains of LZTR1. Similar to SPOP, LZTR1-WT showed punctae immunostaining, whereas the disease-associated LZTR1 BTB-BACK domain mutants which are not able to form multimers and ubiquitinate RAS proteins, show dispersed cytoplasmic localization ^16^. Thus, we speculate that decreased LZTR1 expression due to a loss of one LZTR1 allele might affect oligomer formation, abrogating its ability to ubiquitinate KRAS. Altogether, this suggests that shallow deletion of LZTR1 observed in about 50% of *KRAS*-mutant LUADs results in a full loss of function.

The tumor-suppressive role of LZTR1 is in line with previous studies that reported mutations or deletions of *LZTR1* in various types of cancer ^24^. Our results are supported by recent findings ^41^, showing that the presence of wild-type KRAS in lung adenocarcinomas influences signaling networks. In particular, we found that increased dosage of KRAS-WT protein promoted peritumoral vascular remodeling through activation of the PI3K/mTOR pathway. Previous studies showed that wild-type KRAS activation can induce vascular defects, such as arteriovenous malformations, which are abnormal connections between arteries and veins. KRAS can also promote angiogenesis, which is the formation of new blood vessels from existing ones ^3^. The PI3K-AKT-mTOR-VEGF axis is a key regulator of vascular remodeling, a potential therapeutic target for vascular diseases ^44,45^, and a mediator of therapeutic resistance ^46^.

Furthermore, *Lztr1* loss limits response to the KRAS-G12D inhibitor MRTX113. This result is consistent with recent observations that both mutant-specific and pan-KRAS inhibitors had a limited effect due to the activation of wild-type RAS-like GTPases ^34–36,47^. The feedback activation of wild-type RAS is driven by upstream receptor tyrosine kinases, while pharmacologic inhibition of tyrosine phosphatase SHP2 or epidermal growth factor receptor (EGFR) could abrogate the activation of RAS signaling induced by KRAS-G12C inhibition^34^. However, *LZTR1* depletion leads to resistance to kinase inhibitors, imatinib, and sorafenib as well as to allosteric inhibitors of SHP2 ^14,46,48^, suggesting that co-targeting of KRAS and RTK or SHP2 would not be efficient in cancer cells with *LZTR1* loss. (Re)-activation of LZTR1 activity is more likely to block compensatory activation or secondary mutation of wild-type RAS following the initial inhibition of mutant KRAS ^49^. This report suggests that alternative approaches should be used for tumors with *LZTR1* loss.

mTOR signaling has also emerged as a potential vulnerability in tumors retaining the *KRAS*-WT allele causing resistance to targeted therapy ^41,46^. Targeting PI3K/mTOR signaling has previously been described as a strategy for vascular normalization of tumor vessels in different types of cancers ^45^. This can be exploited, for the design of novel combined therapies targeting both wild-type and mutant KRAS signaling ^50^. The combination therapy of MRTX1133 and dactolisib might confer a risk of toxicity, therefore additional studies will be required to assess the translational potential of our findings. Moreover, we did not test other inhibitors of mutant KRAS, such as Sotorasib, or mTOR inhibitors, such as Everolimus, which may have different pharmacokinetic and pharmacodynamic properties compared to MRTX1133 and dactolisib. In summary, our findings provide a novel therapeutic strategy for lung cancer patients with *KRAS* mutations and *LZTR1* loss.

## Acknowledgments

We thank Prof. Patrick Jacquemin for sharing the YFP-KRAS mice reporters. We thank the KU Leuven Animalium for the animal husbandry and guidance for ethical considerations. We thank Biogenity for performing the proteomics analyses. We thank the University of Iowa Viral Vector Core Facility for the production of adenovirus for *in vivo* injections.

## Author Contribution

TI and RNS performed the *in vivo* experiments; TI and RNS performed the CT scans under the guidance of GVV; RS and GVV analyzed the CT scans; WM performed cloning; TI, WM, and EVB performed the GPS experiments; TI and RNS performed immunoblotting and immunostaining experiments; YM and PZ performed the bioinformatic analyses; ES performed the mathematical modeling; BL performed genotyping and RT-qPCR experiments; TI, RS and AS conceptualized the study, designed the experiments, interpreted the results, and wrote the manuscript; AS acquired financial support for the project. All authors discussed the results and commented on the manuscript.

---

**Supplementary Figure 1:** (A) Immunoblot analysis of RAS expression in single-cell HAP1 clones, in which the GPS cassette for monitoring RAS stability was introduced into the AAVS1 locus. The expression of EGFP-tagged KRAS and pan-RAS were analyzed. (B) Immunoblot analysis of CAS9 expression in single-cell HAP1 clones, in which the GPS cassette was introduced into the AAVS1 locus. (C) Deconvolution of the GPS-based CRISPR screen results. Enrichment of different gRNAs targeting *LZTR1* in sorted vs unsorted GPS reporter HAP1 cells expressing KRAS-WT or KRAS-G12V. Data are shown as Mean±SEM. (D) The DepMap correlation analysis of LZTR1 (x-axis) and KRAS (y-axis) protein expression in lung cancer cell lines harboring wild-type *KRAS* or *KRAS* mutations. Pearson and Spearman scores were calculated for both groups. n=12-13 per group. (E) RT-qPCR analysis of *LZTR*1 mRNA level in lung cancer cell lines expressing sh*GFP* or shRNAs targeting *LZTR1*. Data are shown as MEAN±SEM; n=3. The p-value was calculated using a two-sided t-test.

**Supplementary Figure 2:** Immunostaining for the indicated antibodies on lung tumor sections of *Kras-G12D; Lztr1-WT* and *Kras-G12D; Lztr1-HET* mice 5 weeks and 14 weeks after adenoviral injection with the *Sftpc* specific Cre. Scale bar, 200µm

**Supplementary Figure 3:** (A) Immunoblot analysis of phosphorylated and total MEK1/2 and ERK1/2 in lung cancer isolated from *Kras-G12D; Lztr1-WT* and *Kras-G12D; Lztr1-HET* mice isolated 14 weeks after adenoviral injection with the *Sftpc-*specific Cre. (B) Immunostaining for the indicated antibodies on lung tumor sections of *Kras-G12D; Lztr1-WT* and *Kras-G12D; Lztr1-HET* mice 14 weeks after adenoviral injection with the *Sftpc* specific Cre. Scale bar, 200µm.

## Methods

### Genetically modified mice

All procedures involving animals were performed in accordance with the guidelines of the IACUC of KULeuven and approved project P203/2020. *Lztr1^tm1a^(EUCOMM)^Wtsi^*embryonic stem cells were purchased from EUCOMM (EPD0140_5_E07, EPD0140_5_G06) and used for *in vitro* fertilization of a female CD-1. Primers used for genotyping are:

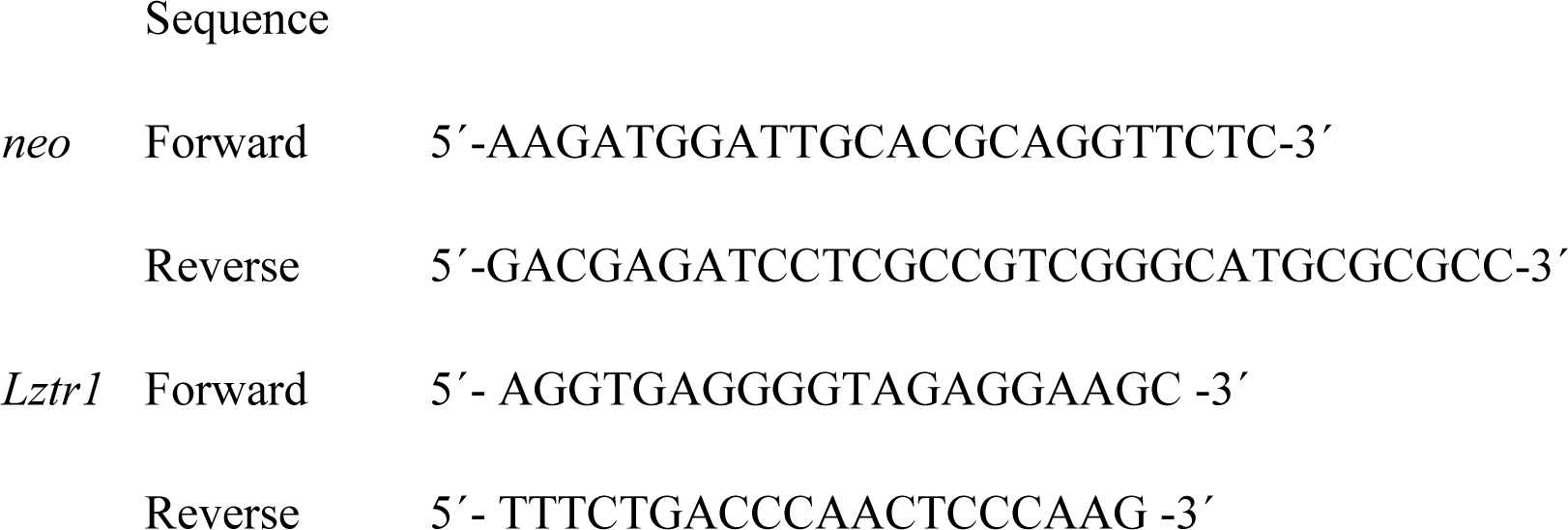

*Lztr1^fl/fl^* mice (*Lztr1^tm1c^(EUCOMM)^Wtsi^*) were generated by Flippase-mediated recombination by breeding *Lztr1^tm1a^(EUCOMM)^Wtsi^* with a *Gt(ROSA)26Sortm1(FLP1)Dym* 1 mice. *Lztr1^fl/fl^* mice were subsequently backcrossed on C57BL6/J background with *B6.129S4-Krastm4Tyj/J (LSL-Kras^G12D^; RRID:IMSR_JAX:008179)* using the project “CreationSablina2020” in accordance with the guidelines of the IACUC of KULeuven. Interbred, *Kras G12D^lsl/wt,^ Lztr1^fl/fl^* mice exhibit normal development, are viable and fertile; and were bred on the “GS2 breeding Sablina” project. The naïve mice showed no disease-related or pain phenotypes. Prof. Anna Sablina is the owner of the mice according to MTAs signed with EUCOMM and with the Jackson Laboratory.

### Plasmids and cloning

The CRISPR-based AAVS1 system for targeted gene insertion into the AAVS1 locus was purchased from Origene (SKU GE100023, SKU GE100024). We cloned the dsRed-IRES-GFP-KRAS-WT-PGK-Hygro and dsRed-IRES-GFP-KRAS-G12V-PGK-Hygro cassettes into the pAAVS1 vector. RAS stability reporter constructs used for transient overexpression were generated using Gateway cloning of the ORFs from corresponding Gateway Entry clones into the MSCV-CMV-DsRed-IRES-EGFP destination vector.

The Gateway plasmids used in this study:

| Plasmid name | Catalog number |
| --- | --- |
| MSCV-CMV-DsRed-IRES-EGFP DEST | Addgene plasmid #41941 |
| pDONR223-KRAS4B | Addgene plasmid #81751 |
| pDONR223-KRAS4B G12V | Addgene plasmid #81665 |
| pDONR223-KRAS4B G12D | Addgene plasmid #81651 |
| pDONR223-KRAS4B G13D | Addgene plasmid #82808 |
| pDONR223-KRAS4B G12C | Addgene plasmid #81667 |
| pDONR223-RIT1 | BCCM Genecorner platform; clone ID: ORFB1045-B03 |
| pDONR223-NRAS | Addgene plasmid #81901 |
| pDONR223-HRAS | Addgene plasmid #81777 |
| pDONR223-MRAS | BCCM Genecorner platform; clone ID: ORFB1102-A06 |

The gRNAs targeting Scramble sequence or *LTR1* locus were introduced in pLenticrispr v2 was a gift from Feng Zhang (Addgene plasmid # 52961).

Sequences:

pLenticrispr-gScramble GCACTACCAGAGCTAACTCA
pLenticrispr-gLZTR1 CTTTACTCAGGGGGTTACAC

shRNA targeting GFP or shRNAs against *LZTR1* was generated by the RNAi Consortium (TRC) and validated previously^1^. pLV-SPB>HA-mKras[NM_001403240.1]:IRES:Cre and pLV-SPB>EGFP:IRES:Cre plasmids were generated by VectorBuilder. All plasmid constructions were verified by DNA sequencing (Eurofins Genomics).

### Cell lines and reagents

EPCAM-positive cells were isolated from mouse lungs after homogenization and digestion with a Multi Tissue Dissociation Kit 2 (Miltenyi Biotech). The digested mix of cells was first incubated with CD45 beads to get rid of CD45 positive cells and then isolated with EPCAM magnetic beads (Miltenyi Biotech). The quality of each fraction was evaluated by immunostaining or immunoblotting for CD45, pan-cytokeratin, and EPCAM.

HAP1 cells (C631) were purchased from Horizon Discovery. H2347 (CRL-5942), SKUL1 (HTB-57), and H727 (CRL-5815) were purchased from ATCC. Cells were cultured in DMEM medium supplemented with 10% FBS (Biowest), 100 µg/ml streptomycin and 100 U/ml penicillin (Gibco). All cell lines were authenticated by STR profiling (ATCC) within the last three years. The experiments were performed with mycoplasma-free cells.

Lentiviral infections were performed as described by the RNAi Consortium (TRC). Infected cells were selected using puromycin for 3 days (2 µg/ml, InvivoGen) or hygromycin B for 7 days (500 µg/ml, InvivoGen). Cells were transfected using GeneJuice (Merck) or Lipofectamine 3000 (ThermoFisher).

HAP1 knock-in RAS stability reporter cells were generated by co-transfection of pCas9-gAAVS1 and pAAVS1-dsRed-IRES-EGFP-KRAS-WT-PGK-Hygro or pAAVS1-dsRed-IRES-EGFP-KRAS-G12V-PGK-Hygro vectors. After hygromycin selection, single cells were sorted using a BD FACS Aria III Cell Sorter (BD Biosciences) and screened by flow cytometry and Sanger sequencing to confirm the incorporation of the reporter cassette into the AAVS1 locus. Single cell clones with the integration of the reporter were transduced with pCas9 and subjected to a second round of single cell sorting (BD FACS Aria III Cell Sorter). For each condition, the clones with a stable reporter profile and the highest Cas9 expression were selected for further experiments.

HEK293T cells were transfected with pLenticrispr-g*LZTR1* or pLenticrispr-g*Scramble* as a control. 24h post transfection, cells were subjected to puromycin selection. Genomic DNA was extracted using a Nucleospin tissue kit (Machéry-Nagel, 7409522) and *LZTR1*-specific primers were used to amplify the region around the cutting site (FW: CTCCACCTTCCAGGGTTTGAA and RV: ATGGCTACATCCACCCGACA). *LZTR1* indel was then verified by Sanger sequencing using the *LZTR1*-specific primers.

Drugs were purchased from MedChemExpress and used at the dose indicated below:

|  | *In vitro* | *In vivo* |
| --- | --- | --- |
| MG132 | 10 µM | - |
| MRTX1133 | 1 to 200nM | 10mg/kg |
| Dactolisib (NVP BEZ 235) | 1 μM | 10mg/kg |
| Everolimus | 1 μM | - |
| Ipatasertib | 1 μM | - |
| Selumetinib | 1 μM | - |
| Trametinib | 1 μM | - |

### Protein stability assay

24 hours after plating, HEK293T cells were transfected with the RAS stability reporter plasmids. 48 hours after transfection, cells were harvested and analysed using the MACSQuant VYB Flow Cytometer (Miltenyi Biotec). Raw data were analysed using FlowJo Software (BD Biosciences). Live single cells were monitored for the expression of EGFP and dsRed, and the EGFP/DsRed ratio was counted to measure the relative protein stability.

### FACS-based genome-wide CRISPR screen of RAS stability modifiers

HAP1 knock-in RAS stability reporter (wt-KRAS or KRAS-G12V) cells were infected with a lentiviral pool of the Human Brunello CRISPR knockout pooled library (a gift from David Root and John Doench: Addgene #73178) at an MOI of 0.3 and with a coverage of 500 cells per sgRNA. 24h after transduction, the cells were washed with PBS; and fresh growth medium supplemented with 2µg/ml puromycin was added to the cells. After 2 days of puromycin selection, the medium was replaced with growth medium without puromycin. 5 days after transduction, cells were harvested and sorted on a BD FACS Aria III or BD FACSAria™ Fusion Cell Sorter (BD Biosciences). Both unsorted population and the top 2% of cells with the highest EGFP/dsRed ratio were collected. The screen was performed in triplicates.

DNA extraction was performed on collected populations using a Nucleospin tissue kit (Machéry-Nagel, 7409522) according to the manufacturer’s instructions. To prepare the sgRNA library for sequencing, the sgRNA regions were amplified from the genomic DNA by PCR using the NEBNext Q5 HotStart HiFi polymerase (NEB, M0543) and specific primers. Amplified sequences were purified with a QiaQuick PCR purification kit (Qiagen, 28104). Further library preparation and Illumina sequencing (NovaSeq PE150) was performed by Novogene.

For the CRISPR screen analysis, as the gRNA is located at position 31 in each read, as specified by the sequencing protocol, we used a custom Perl script to directly extract gRNA sequences from the FASTQ reads. We measured the abundance of each gRNA sequence by counting the number of reads aligned to it. To ensure reliable analysis, we kept only gRNAs having a median of at least 10 reads across all samples. The read counts of remaining gRNAs were normalized to reads per million (RPM) to adjust for the differences in sequencing depth. After normalization, RPM values were converted by log2. To minimize batch effects, we first compared each replicate independently. Following this, to measure the effect of gRNAs within each comparison, we calculated the difference between the log2 transformed RPM values for each gRNA between unsorted and sorted cells. Next, we multiplied this difference by the sum of the log2 transformed RPM values in both conditions for each gRNA, which emphasizes the effect of gRNAs with higher total abundance across conditions. Finally, we multiplied this value by the rank of the log2 difference in the list of all gRNAs, which highlights gRNAs with the most significant impact relative to others. To obtain a representative value for each gRNA, we computed the average across the replicates. To assess the overall effect of each gene, the representative values obtained from all gRNAs targeting the same gene were averaged again.

### Intratracheal Adenoviral injection

The Adenovirus expressing Cre recombinase under Surfactant Protein C (*Sftpc*) promoter was purchased from The University of Iowa Gene Transfer Vector Core (https://vector-core.medicine.uiowa.edu/collections/ad5/products/ad5mspc-cre). The virus was added to MEM and prepared in a sterile syringe with a 23-20-gauge tracheal tubing or a 22-gauge venous catheter. The injected volume was around 0.05 ml. The mouse was anesthetized with an appropriate method, such as intraperitoneal injection of ketamine. The mouse was placed in the prone position on a surgery platform. The neck region of the mouse was exposed and disinfected with povidone-iodine. A small incision was made in the skin and subcutaneous tissue to expose the trachea. Alternatively, a non-invasive method that did not require an incision was used, such as inserting the tubing through the mouth and into the trachea. The tubing was carefully inserted into the trachea, avoiding damage to the surrounding tissues. The substance was slowly injected into the trachea, ensuring that there was no leakage or reflux. The chest movement of the mouse was monitored to confirm that the substance reached the lungs. The tubing was withdrawn, and the incision was closed with sutures. The mouse was monitored until it recovered from anesthesia and provided analgesia if needed.

### Computed micro-computed tomography (micro-CT)

Before the start of the imaging sessions animals underwent at least 1 week of acclimatization to the new environment. Animals were anesthetized using isoflurane (induction at 3% in 100% oxygen and maintenance at 1-2% in 100% oxygen). The animals were scanned on a dedicated in vivo micro-CT scanner (SkyScan 1278, Bruker micro-CT, Kontich, Belgium) at an X-ray dose that is well tolerated ^2–4^. The images were acquired with the following parameters: 65 kVp X-ray source voltage and 350 μA current combined with an X-ray filter of 1 mm aluminum, 150 ms exposure time per projection, 3 projections averaged per view, acquiring projections with 0.9° increments over a total angle of 220°, producing reconstructed 3D data sets with 50 μm isotropic reconstructed voxel size. The total scanning time per mouse was approximately 2.5 min, resulting in a measured delivered radiation dose of approximately 60-80 mGy per scan. A baseline scan was performed before the adenoviral injection, and 4 weeks after injection the scans were performed every two weeks. The total nodules volume was evaluated after segmentation of the lung using MIPAV (NIH). Next, each region is analyzed to extract features, such as shape, texture, and intensity, that can help identify the nodules. Then, the nodules are separated from the blood vessels using a mathematical formula called the sphericity parameter, which measures how spherical an object is. The more spherical an object is, the more likely it is a nodule and not a vessel. Finally, the volume of each nodule is calculated using a technique called volumetric analysis, which estimates the volume occupied by nodules, using the volumetric neuroimage analysis extension of MIPAV, according to the methodology described before^5^.

### Optical projection tomography of cleared lungs

The clearing of lungs was performed according to Life Canvas guidelines, using the passive clearing protocol for the Passive Clearing Kit (C-PCK-250). Immunostaining for Smooth Muscle Actin (SMA) was performed according to the guidelines of Life Canvas. The cleared lungs were scanned on a dedicated Optical projection tomography Bioptonics 3001 (Skyscan, Kontich, Belgium). The images were acquired with the following parameters: TexasRed laser power, excitation 560/40nm, emission 610nm, 100 ms exposure time per projection, 3 projections averaged per view, acquiring projections with 0.9° increments over a total angle of 360°, producing reconstructed 3D data sets with 6.7 μm isotropic reconstructed voxel size. Quantification of vascular tortuosity on volumes after reconstruction of the vascular network using Bitplane Imaris Surpass mode, skeletonization of the network using Skeletonize3D plugin, and quantification of morphometric parameters using the AnalyzeSkeleton plugin, according to approaches validated previously^6,7^.

### Cytokine array and ELISA assay

The cytokine array to detect mouse and human cytokines was performed according to the manufacturers’ protocol (Raybiotech). The array was scanned using the TECCAN Powerscanner and analysed with the software provided. Mouse Cytokine Array C1 (AAM-CYT-1) (Raybiotech). ELISA assay to measure VEGFA levels was performed according to the manufacturers’ protocol (Biolegend, 430204) using the lung cancer cell supernatant.

### Antibodies

The following antibodies/dyes were used:

|  |  |
| --- | --- |
| Mouse monoclonal | Anti-Vinculin (Sigma Aldrich, clone hVIN-1, V 9131), anti β-actin (Sigma-Aldrich, A5441), anti Cas9 (sc-517386, SCBT), Anti EIF4E (P85D11B4, Biolegend), anti LZTR1 (E-12) (sc-390166, SCBT), anti mTOR (6H9B10, Biolegend), anti phospho-mTOR (S2448) (A17024A, Biolegend), anti KRAS-G12D mutant (GTX635362, GeneTex), anti KRAS-G12V mutant (GTX635623, GeneTex), Anti-RIT1 (GTX117329, GeneTex), |
| Rabbit polyclonal | Anti AKT1 (#2938, Cell Signaling), anti Phospho-AKT1 (S473) (#9018, Cell Signaling), anti Cytokeratin 19 (10712-1-AP, Proteintech), anti phospho EIF4E (S209) (#9741T, Cell Signaling), anti FLAG Epitope (DYKDDDDK) (740001, ThermoFisher), anti GATA6 (55435-1-AP, Proteintech), anti GFP (NB600-308, Novus Biologicals), anti HIF1α (H1alpha67, Novus Biologicals), anti KRAS (12063-1-AP, Proteintech), anti phospho-MEK1/2 (S217/S221) (#9121, Cell Signaling), anti MEK1/2 (#9122S, Cell Signaling), anti pan-RAS (MABS195, EMD Millipore), p44/42 MAPK (ERK1/2) (#9102, Cell Signaling), anti phospho-p44/42 MAPK (ERK1/2) (T202/Y204) (#4370, Cell Signaling), anti TTF1 (66034-1-Ig, Proteintech). |
| Rat monoclonal | Anti-CD31 (Ms,Sw) from Rat (DIA310, Dianova) |
| Coupled antibodies | Anti-β-actin, alpha-Smooth Muscle - Cy3 antibody (C6198 – MilliporeSigma), Alexa Fluor® 488 anti-mouse CD54 (ICAM1) Antibody (116112, BioLegend), Brilliant Violet 605™ anti-mouse/human CD11b (101237, BioLegend) |

The following secondary antibodies were used: Donkey or Goat Alexa488-or Alexa568 or Alexa594-or Alexa 647-conjugated secondary antibodies (ThermoFisher, Molecular Probes) (A10037, A10042, A11005, A11006, A11012, A11034, A11057, A21201, A21245, A21247, A21447) or HRP-labelled antibodies (DAKO) (P0447, P0448).

### Immunoblotting

Cells were washed in cold PBS and scraped on ice in NP40 lysis buffer (50 mM Tris-HCl pH 7.5, 150 mM NaCl, 1% NP40, 5% glycerol) or RIPA lysis buffer (50 mM Tris-HCl pH 7.5, 150 mM NaCl, 1% NP40, 0.5% sodium deoxycholate, 0.1% SDS) containing protease inhibitor cocktail (Merk Millipore) and phosphatase inhibitors (Sigma-Aldrich). Cell lysates were cleared for 15 minutes at 16 000 x g at 4°C. Protein concentration was quantified using a Pierce BCA protein assay kit (ThermoFisher). For immunoblotting, proteins were separated using SDS-PAGE gels (ThermoFisher) and transferred to nitrocellulose membranes. Membranes were immunoblotted with the primary and secondary HRP-conjugated antibodies. The chemiluminescence was detected with the chemiluminescent substrate kit (ThermoFisher) using the digital developer.

### Immunocytochemistry and immunohistochemistry

For immunocytochemistry, 2 x 10^4^ cells per well were seeded on an 8-well chamber glass slide (Ibidi). Cells were fixed with 4% paraformaldehyde in PBS for 15 minutes at room temperature. Cells were permeabilized with 0.1% Triton-X100 in PBS and blocked with 1% goat serum for 1 hour at room temperature. Primary antibodies and goat fluo-conjugated secondary antibodies were diluted in 1% goat serum-blocking buffer. Cells were mounted using Vectashield antifade mounting medium with DAPI (Vector Laboratories).

For immunohistochemistry, tumors were fixed in 4% paraformaldehyde and embedded in paraffin. Paraffin slides were rehydrated and treated with hydrogen peroxide. Antigen retrieval was performed by heat in Tris-EDTA pH 9.0. The sections were incubated with primary and secondary antibodies, and diaminobenzidine (Dako) was used as a detection method followed by hematoxylin counterstaining. Quantification of the IHC immunostainings was done using the ImageJ IHC Profiler plugin.

### Microscopic analysis

Microscopic analysis was done with a Leica DCF-6000 confocal microscope; images acquired with LAS X software. Some images were acquired using a Nikon SMZ25 stereo zoom microscope (Perfect zoom 3.15-315x) and acquired using ZEN Blue (ZEISS). Image analysis was performed with Imaris (Bitplane), Zeiss Aviovision or Image J. Imaged fields for quantification were randomly selected on the tissue section or cell monolayer. Representative images were chosen after the quantification, as the most accurate depiction of the phenotype of the control when compared to the experimental condition.

### Mass Spectrometry Sample Digestion

The cell pellets and tissue were lysed with lysis buffer consisting of 6M Guanidinium Hydrochloride, 10 mM TCEP, 40 mM CAA, 50 mM HEPES pH8.5. Samples were boiled at 95°C for 5 minutes, processed using the TissueLyser for 2 times for 1min going from 3 to 30hz and sonicated on high for 5x 60 seconds on and 30 seconds off in a Bioruptor Pico sonication water bath (Diagenode) at 4°C. Sample concentrations were determined using BCA and 200µg of sample were taken forward for digestion. Samples were diluted 1:3 with 10% Acetonitrile, 50 mM HEPES pH 8.5, LysC (MS grade, Wako) was added in a 1:50 (enzyme to protein) ratio, and samples were incubated at 37°C for 4hrs. Samples were further diluted to a final 1:10 with 10% Acetonitrile, 50 mM HEPES pH 8.5, trypsin (MS grade, Sigma) was added in a 1:100 (enzyme to protein) ratio and samples were incubated overnight at 37°C. Enzyme activity was quenched by adding 2% trifluoroacetic acid (TFA) to a final concentration of 1%. Prior to further processing, the peptides were desalted on a SOLAµ SPE plate (HRP, Thermo). For each sample, the filters were activated with 200ul of 100% Methanol (HPLC grade, Sigma), then 200ul of 80% Acetonitrile, 0.1% formic acid. The filters were subsequently equilibrated 2x with 200ul of 1% TFA, 3% Acetonitrile, after which the sample was loaded using centrifugation at 1,500x rpm. After washing the tips twice with 200ul of 0.1% formic acid, the peptides were eluted into clean 1.5ml Eppendorf tubes using 40% Acetonitrile, 0.1% formic acid. The eluted peptides were concentrated in an Eppendorf Speedvac and re-constituted in 50mM HEPES (pH8.5).

### TMT quantitative proteomics analysis

For TMT labeling with 16plex tags (ThermoFischer, TFA was added to the samples to bring acetonitrile concentration down to less than 5%. Prior to mass spectrometry analysis, the peptides were fractionated using an offline ThermoFisher Ultimate3000 liquid chromatography system using high pH fractionation (5mM Ammonium Bicarbonate, pH 10) at 5ul/min flowrate. 15µg of peptides were separated over a 120min gradient (5% to 35% Acetonitrile), while collecting fractions every 130sec. The resulting 60 fractions were pooled into 30 final fractions, acidified to pH < 2 with 1% TFA and loaded onto EvoSep stagetips according to manufacturer’s protocol.

For each fraction, peptides were analyzed using the pre-set ‘30 samples per day’ method on the EvoSep One instrument. Peptides were eluted over a 44-min gradient and analysed with an Orbitrap Eclipse^TM^ Tribrid^TM^ instrument (Thermo Fisher) with FAIMS Pro^TM^ Interface (ThermoFisher) switched between CVs of −50 V and −70 V with cycle times of 2 s and 1.5 s respectively. Full MS spectra were collected at a resolution of 120,000, with normalized AGC target set to 100% or maximum injection time of 50 ms and a scan range of 375–1500 m/z. MS1 precursors with an intensity of >5×10^3^ and charge state of 2-7 were selected for MS2 analysis. Dynamic exclusion was set to 120 s, the exclusion list was shared between CV values and Advanced Peak Determination was set to ‘off’. The precursor fit threshold was set to 70% with a fit window of 0.7 m/z for MS2. Precursors selected for MS2 were isolated in the quadrupole with a 0.7 m/z window. Ions were collected for a maximum injection time of 35 ms and normalized AGC target set to 300%. Fragmentation was performed with a HCD normalized collision energy of 30% and MS2 spectra were acquired in the IT at scan rate rapid. The MS2 spectra were subjected to RTS using the Uniprot protein database Homo sapiens (and trypsin set as enzyme. Static modifications were TMTpro on Lysine (K) and N-terminus and carbamidomethyl on Cysteine (C). Oxidation of methionine (M) was set as variable modification. The Maximum missed cleavages parameter was set to 1 and maximum variable modifications to 2. FDR filtering enabled, maximum search time was set to 35 ms, and the scoring threshold was set to 1 Xcorr, 0 dCn, and 5 ppm precursor tolerance. Use as trigger only was disabled and close-out was enabled with maximum number of peptides per protein set to 4. Precursors were subsequently filtered with an isobaric tag loss exclusion of TMT and precursor mass exclusion set to25 ppm low and 25 ppm high. Precursors identified by RTS were isolated for an MS3 scan using the quadrupole with a 2 m/z window, and ions were collected for a maximum injection time of 86 ms and normalized AGC target of 300%. Turbo TMT was deactivated, and the number of dependent scans set to 10. Isolated precursors were fragmented again with 50% normalized HCD collision energy, and MS3 spectra were acquired in the orbitrap at 50000 resolution with a scan range of 100-500 m/z. MS performance was verified for consistency by running complex cell lysate quality control standard.

The raw files were analyzed using Proteome Discoverer 2.4 (ThermoFisher). TMT reporter ion quantitation was enabled in the processing and consensus steps, and spectra were matched against the Uniprot Homo sapiens database, including reviewed and unreviewed proteins. Dynamic modifications were set as Oxidation (M) and Acetyl on protein N-termini. Cysteine carbamidomethyl (on C residues) and TMT 16-plex (on peptide N-termini and K residues) were set as static modifications. All results were filtered to a 1% FDR, and protein quantitation done using the built-in Minora Feature Detector with statistical significance testing done with the built-in t-test.

### Label-free quantitative phosphoproteomic analysis

Phosphopeptides were enriched using MagResyn Ti-IMAC beads (LabLife). 25µL of bead particles were first washed with 200µL 70% ethanol, then in 100µL 1% NH4OH, and finally washed three times in a solution of 80% Acetonitrile, 1M glycolic acid and 5% TFA (loading buffer). Samples were mixed in a 1:1 ratio with this same loading buffer, before being incubated with the beads for 30min at RT. Supernatant was removed and beads were wash in 400µL of loading buffer. Afterwards, beads were washed twice in 400µL 80% ACN with 1% TFA and twice in 400µL 10% ACN with 0.1% TFA. After the final wash, beads were transferred to clean Eppendorf tubes and incubated 3 times in 80µL 1% NH_4_OH to elute peptides from beads. The eluted samples were speedvac’ed for 30mins at 60C before being acidified and desalted on SOLAµ SPE plate (HRP, ThermoFischer), following the same procedure as previously. Dried peptides were reconstituted in 12µL 2% ACN, 1%TFA/20µL HEPES 50mM pH 8.5.

Peptides were loaded onto a 2cm C18 trap column (ThermoFisher 164946), connected in-line to a 15cm C18 reverse-phase analytical column (Thermo EasySpray ES904) using 100% Buffer A (0.1% Formic acid in water) at 750bar, using the Thermo EasyLC 1200 HPLC system, and the column oven operating at 30°C. Peptides were eluted over a 70 minute gradient ranging from 10% to 60% of 80% acetonitrile, 0.1% formic acid at 250 nl/min, and the Orbitrap Eclipse^TM^ Tribrid^TM^ instrument (ThermoFisher) was run in DDA mode with FAIMS Pro^TM^ Interface (ThermoFisher) switched between CVs of −50 V and −70 V with cycle times of 2 s and 1.5 s respectively. Full MS spectra were collected at a resolution of 120,000, with an AGC target of 100% or maximum injection time set to ‘auto’ and a scan range of 375–1500 m/z. The MS2 spectra were obtained in the orbitrap operating at a resolution of 60.000, with an AGC target of 100% or maximum injection time set to ‘auto’, a normalized HCD collision energy of 30 and an intensity threshold of 2.5e4. Dynamic exclusion was set to 60 s, and ions with a charge state <2, >7 or unknown were excluded. MS performance was verified for consistency by running complex cell lysate quality control standards, and chromatography was monitored to check for reproducibility.

The raw files were analyzed using Proteome Discoverer 2.4. Label-free quantitation (LFQ) was enabled in the processing and consensus steps, and spectra were matched against the Uniprot Homo sapiens database, including reviewed and unreviewed proteins. Dynamic modifications were set as Oxidation (M), Phospho(S, T, Y) and Acetyl on protein N-termini. Cysteine carbamidomethyl was set as a static modification. All results were filtered to a 1% FDR, and protein quantitation done using the built-in Minora Feature Detector.

### Gene expression analysis of the TCGA-LUAD cohort

Counts files for TCGA-LUAD samples were acquired from GDC portal (https://portal.gdc.cancer.gov/projects/TCGA-LUAD). Only *KRAS* mutant patients, identified using WGS data from TCGA Pancancer Atlas, were included in subsequent analyses. Patients were classified according *LZTR1* copy number variation status as *LZTR1-*WT (diploid) or *LZTR1-*HETLOSS (shallow deletion) based on WGS information obtained from *cBioPortal*. After conducting a differential gene expression analysis using DEseq2, a pre-ranked GSEA analysis was performed.

### mRNA expression analysis

RNA was isolated using the NucleoSpin RNA kit (Machery Nagel). 500 ng of RNA was used for reverse transcription with a Sensifast cDNA synthesis kit (Bioline). RT-qPCR was performed with LightCycler 480 SYBR Green I Master reagent with the primers listed below.

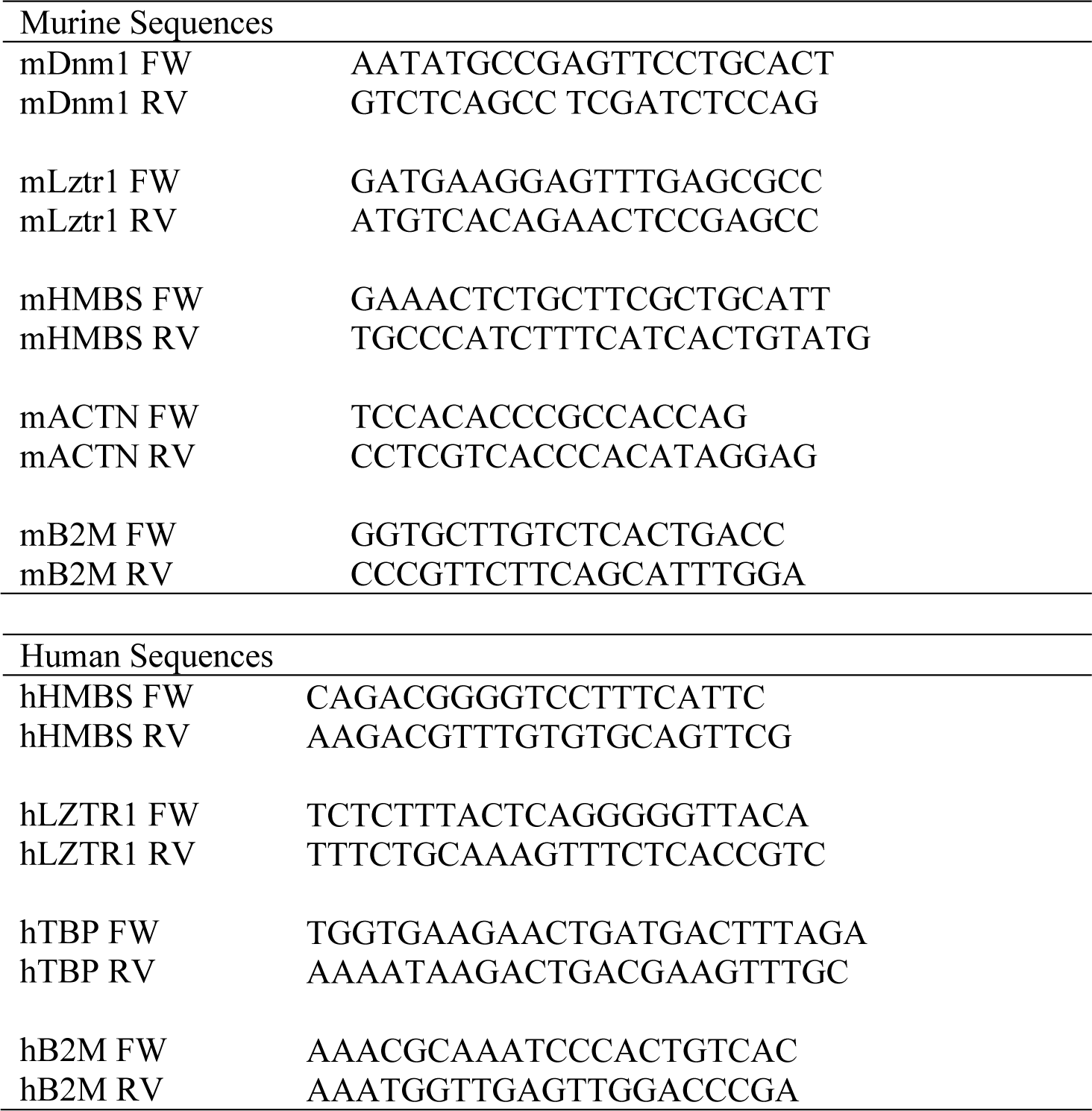
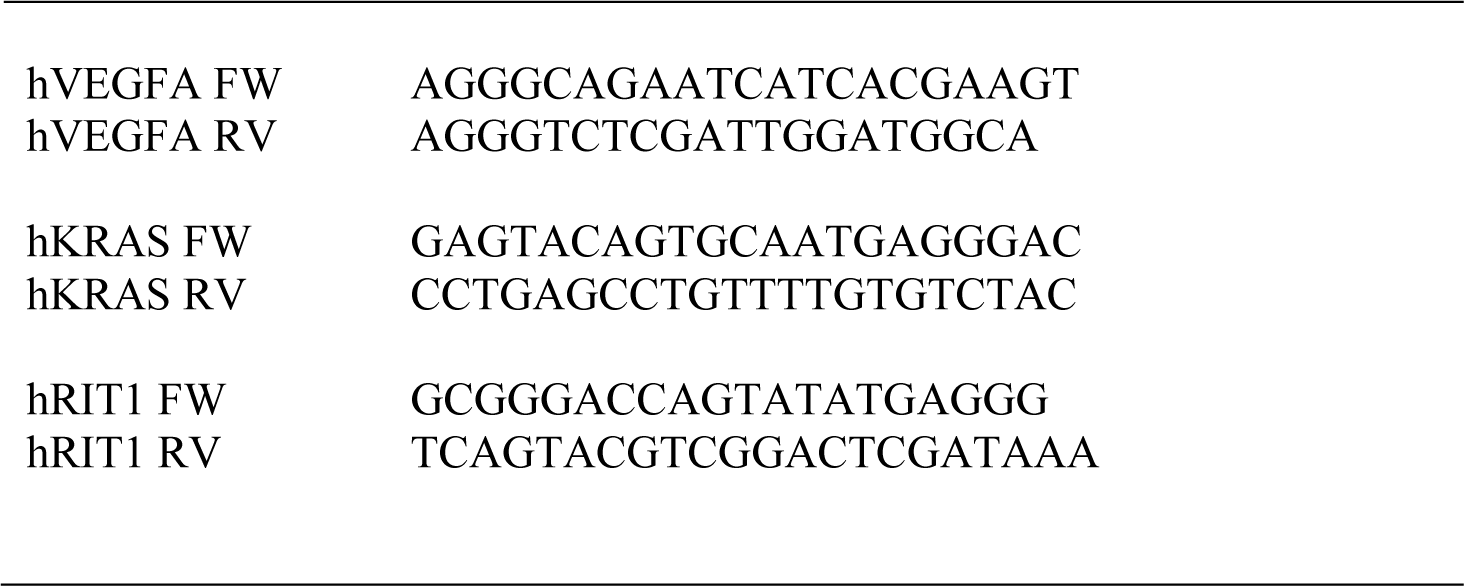

### Mathematical modeling of RAS signaling

The mathematical model of RAS signaling, as regulated by RAS GEFs, RAS GAPs, intrinsic GTPase activity, spontaneous nucleotide dissociation, spontaneous nucleotide association, and effort binding/unbinding has previously been developed and described in detail ^8,9^. For estimates here, the base parameter set of protein abundances, enzymatic activities, and KRAS G12D parameters was used ^9^. The model estimates that 50% of total RAS is KRAS and the other 50% is split between NRAS and HRAS, and models half of the KRAS population with wild-type parameters and half with mutant parameters ^8,10^.

### Statistics

Data entry and all analyses were performed blindly. All statistical analyses were performed using GraphPad Prism software assuming non-parametric parameters. Statistical significance was calculated by Wilcoxon Mann-Whitney or Wilcoxon matched-pairs signed-rank on two experimental conditions. A comparison of continuous variables between more than two groups was performed by Kruskal-Wallis test and if statistical significance was observed, Dunn’s Multiple Comparison Test. Two-dimensional data were analyzed using mixed model analysis, using Graphpad PRISM version 9. Graphs show each replicate as dots. No experiment-wide multiple-test correction was applied. Proteomic and phosphoproteomic data were analyzed using Ingenuity pathway analysis (IPA) (QIAGEN Inc).

## References

1. Cancer Genome Atlas Research, N. (2014). Comprehensive molecular profiling of lung adenocarcinoma. Nature 511, 543–550. 10.1038/nature13385.

2. Colicelli, J. (2004). Human RAS superfamily proteins and related GTPases. Sci STKE 2004, RE13. 10.1126/stke.2502004re13.

3. Simanshu, D.K., Nissley, D.V., and McCormick, F. (2017). RAS Proteins and Their Regulators in Human Disease. Cell 170, 17–33. 10.1016/j.cell.2017.06.009.

4. Burgess, M.R., Hwang, E., Mroue, R., Bielski, C.M., Wandler, A.M., Huang, B.J., Firestone, A.J., Young, A., Lacap, J.A., Crocker, L., et al. (2017). KRAS Allelic Imbalance Enhances Fitness and Modulates MAP Kinase Dependence in Cancer. Cell 168, 817–829 e815. 10.1016/j.cell.2017.01.020.

5. Grabocka, E., Pylayeva-Gupta, Y., Jones, M.J., Lubkov, V., Yemanaberhan, E., Taylor, L., Jeng, H.H., and Bar-Sagi, D. (2014). Wild-type H- and N-Ras promote mutant K-Ras-driven tumorigenesis by modulating the DNA damage response. Cancer Cell 25, 243–256. 10.1016/j.ccr.2014.01.005.

6. Tang, R., Shuldiner, E.G., Kelly, M., Murray, C.W., Hebert, J.D., Andrejka, L., Tsai, M.K., Hughes, N.W., Parker, M.I., Cai, H., et al. (2023). Multiplexed screens identify RAS paralogues HRAS and NRAS as suppressors of KRAS-driven lung cancer growth. Nat Cell Biol 25, 159–169. 10.1038/s41556-022-01049-w.

7. Weyandt, J.D., Lampson, B.L., Tang, S., Mastrodomenico, M., Cardona, D.M., and Counter, C.M. (2015). Wild-Type Hras Suppresses the Earliest Stages of Tumorigenesis in a Genetically Engineered Mouse Model of Pancreatic Cancer. PLoS One 10, e0140253. 10.1371/journal.pone.0140253.

8. Young, A., Lou, D., and McCormick, F. (2013). Oncogenic and wild-type Ras play divergent roles in the regulation of mitogen-activated protein kinase signaling. Cancer Discov 3, 112–123. 10.1158/2159-8290.CD-12-0231.

9. Lo, A., Holmes, K., Kamlapurkar, S., Mundt, F., Moorthi, S., Fung, I., Fereshetian, S., Watson, J., Carr, S.A., Mertins, P., and Berger, A.H. (2021). Multiomic characterization of oncogenic signaling mediated by wild-type and mutant RIT1. Sci Signal 14, eabc4520. 10.1126/scisignal.abc4520.

10. Cuevas-Navarro, A., Wagner, M., Van, R., Swain, M., Mo, S., Columbus, J., Allison, M.R., Cheng, A., Messing, S., Turbyville, T.J., et al. (2023). RAS-dependent RAF-MAPK hyperactivation by pathogenic RIT1 is a therapeutic target in Noonan syndrome-associated cardiac hypertrophy. Sci Adv 9, eadf4766. 10.1126/sciadv.adf4766.

11. Young, L.C., and Rodriguez-Viciana, P. (2018). MRAS: A Close but Understudied Member of the RAS Family. Cold Spring Harb Perspect Med 8. 10.1101/cshperspect.a033621.

12. Bahar, M.E., Kim, H.J., and Kim, D.R. (2023). Targeting the RAS/RAF/MAPK pathway for cancer therapy: from mechanism to clinical studies. Signal Transduct Target Ther 8, 455. 10.1038/s41392-023-01705-z.

13. Abe, T., Umeki, I., Kanno, S.I., Inoue, S.I., Niihori, T., and Aoki, Y. (2020). LZTR1 facilitates polyubiquitination and degradation of RAS-GTPases. Cell Death Differ 27, 1023–1035. 10.1038/s41418-019-0395-5.

14. Bigenzahn, J.W., Collu, G.M., Kartnig, F., Pieraks, M., Vladimer, G.I., Heinz, L.X., Sedlyarov, V., Schischlik, F., Fauster, A., Rebsamen, M., et al. (2018). LZTR1 is a regulator of RAS ubiquitination and signaling. Science 362, 1171–1177. 10.1126/science.aap8210.

15. Castel, P., Cheng, A., Cuevas-Navarro, A., Everman, D.B., Papageorge, A.G., Simanshu, D.K., Tankka, A., Galeas, J., Urisman, A., and McCormick, F. (2019). RIT1 oncoproteins escape LZTR1-mediated proteolysis. Science 363, 1226–1230. 10.1126/science.aav1444.

16. Steklov, M., Pandolfi, S., Baietti, M.F., Batiuk, A., Carai, P., Najm, P., Zhang, M., Jang, H., Renzi, F., Cai, Y., et al. (2018). Mutations in LZTR1 drive human disease by dysregulating RAS ubiquitination. Science 362, 1177–1182. 10.1126/science.aap7607.

17. Paganini, I., Chang, V.Y., Capone, G.L., Vitte, J., Benelli, M., Barbetti, L., Sestini, R., Trevisson, E., Hulsebos, T.J., Giovannini, M., et al. (2015). Expanding the mutational spectrum of LZTR1 in schwannomatosis. Eur J Hum Genet 23, 963–968. 10.1038/ejhg.2014.220.

18. Piotrowski, A., Xie, J., Liu, Y.F., Poplawski, A.B., Gomes, A.R., Madanecki, P., Fu, C., Crowley, M.R., Crossman, D.K., Armstrong, L., et al. (2014). Germline loss-of-function mutations in LZTR1 predispose to an inherited disorder of multiple schwannomas. Nat Genet 46, 182–187. 10.1038/ng.2855.

19. Yamamoto, G.L., Aguena, M., Gos, M., Hung, C., Pilch, J., Fahiminiya, S., Abramowicz, A., Cristian, I., Buscarilli, M., Naslavsky, M.S., et al. (2015). Rare variants in SOS2 and LZTR1 are associated with Noonan syndrome. J Med Genet 52, 413–421. 10.1136/jmedgenet-2015-103018.

20. Mermel, C.H., Schumacher, S.E., Hill, B., Meyerson, M.L., Beroukhim, R., and Getz, G. (2011). GISTIC2.0 facilitates sensitive and confident localization of the targets of focal somatic copy-number alteration in human cancers. Genome Biol 12, R41. 10.1186/gb-2011-12-4-r41.

21. Inoue, D., Polaski, J.T., Taylor, J., Castel, P., Chen, S., Kobayashi, S., Hogg, S.J., Hayashi, Y., Pineda, J.M.B., El Marabti, E., et al. (2021). Minor intron retention drives clonal hematopoietic disorders and diverse cancer predisposition. Nat Genet 53, 707–718. 10.1038/s41588-021-00828-9.

22. Sewduth, R.N., Pandolfi, S., Steklov, M., Sheryazdanova, A., Zhao, P., Criem, N., Baietti, M.F., Lechat, B., Quarck, R., Impens, F., and Sablina, A.A. (2020). The Noonan Syndrome Gene Lztr1 Controls Cardiovascular Function by Regulating Vesicular Trafficking. Circ Res 126, 1379–1393. 10.1161/CIRCRESAHA.119.315730.

23. Ivanisevic, T., Steklov, M., Lechat, B., Cawthorne, C., Gsell, W., Velde, G.V., Deroose, C., Van Laere, K., Himmelreich, U., Sewduth, R.N., and Sablina, A.A. (2023). Targeted STAT1 therapy for LZTR1-driven peripheral nerve sheath tumor. Cancer Commun (Lond) 43, 1386–1390. 10.1002/cac2.12490.

24. Ko, A., Hasanain, M., Oh, Y.T., D’Angelo, F., Sommer, D., Frangaj, B., Tran, S., Bielle, F., Pollo, B., Paterra, R., et al. (2023). LZTR1 Mutation Mediates Oncogenesis through Stabilization of EGFR and AXL. Cancer Discov 13, 702–723. 10.1158/2159-8290.CD-22-0376.

25. Abe, T., Umeki, I., Kanno, S.I., Inoue, S.I., Niihori, T., and Aoki, Y. (2019). LZTR1 facilitates polyubiquitination and degradation of RAS-GTPases. Cell Death Differ. 10.1038/s41418-019-0395-5.

26. Yen, H.C., and Elledge, S.J. (2008). Identification of SCF ubiquitin ligase substrates by global protein stability profiling. Science 322, 923–929. 10.1126/science.1160462.

27. Assi, M., Achouri, Y., Loriot, A., Dauguet, N., Dahou, H., Baldan, J., Libert, M., Fain, J.S., Guerra, C., Bouwens, L., et al. (2021). Dynamic Regulation of Expression of KRAS and Its Effectors Determines the Ability to Initiate Tumorigenesis in Pancreatic Acinar Cells. Cancer Res 81, 2679–2689. 10.1158/0008-5472.CAN-20-2976.

28. Kovalski, J.R., Bhaduri, A., Zehnder, A.M., Neela, P.H., Che, Y., Wozniak, G.G., and Khavari, P.A. (2019). The Functional Proximal Proteome of Oncogenic Ras Includes mTORC2. Mol Cell 73, 830–844 e812. 10.1016/j.molcel.2018.12.001.

29. Timms, R.T., Mena, E.L., Leng, Y., Li, M.Z., Tchasovnikarova, I.A., Koren, I., and Elledge, S.J. (2023). Defining E3 ligase-substrate relationships through multiplex CRISPR screening. Nat Cell Biol 25, 1535–1545. 10.1038/s41556-023-01229-2.

30. Li, S., Liu, S., Deng, J., Akbay, E.A., Hai, J., Ambrogio, C., Zhang, L., Zhou, F., Jenkins, R.W., Adeegbe, D.O., et al. (2018). Assessing Therapeutic Efficacy of MEK Inhibition in a KRAS(G12C)-Driven Mouse Model of Lung Cancer. Clin Cancer Res 24, 4854–4864. 10.1158/1078-0432.CCR-17-3438.

31. McFall, T., Diedrich, J.K., Mengistu, M., Littlechild, S.L., Paskvan, K.V., Sisk-Hackworth, L., Moresco, J.J., Shaw, A.S., and Stites, E.C. (2019). A systems mechanism for KRAS mutant allele-specific responses to targeted therapy. Sci Signal 12. 10.1126/scisignal.aaw8288.

32. McFall, T., and Stites, E.C. (2021). Identification of RAS mutant biomarkers for EGFR inhibitor sensitivity using a systems biochemical approach. Cell Rep 37, 110096. 10.1016/j.celrep.2021.110096.

33. Stites, E.C., Trampont, P.C., Ma, Z., and Ravichandran, K.S. (2007). Network analysis of oncogenic Ras activation in cancer. Science 318, 463–467. 10.1126/science.1144642.

34. Ryan, M.B., Coker, O., Sorokin, A., Fella, K., Barnes, H., Wong, E., Kanikarla, P., Gao, F., Zhang, Y., Zhou, L., et al. (2022). KRAS(G12C)-independent feedback activation of wild-type RAS constrains KRAS(G12C) inhibitor efficacy. Cell Rep 39, 110993. 10.1016/j.celrep.2022.110993.

35. Kim, D., Herdeis, L., Rudolph, D., Zhao, Y., Bottcher, J., Vides, A., Ayala-Santos, C.I., Pourfarjam, Y., Cuevas-Navarro, A., Xue, J.Y., et al. (2023). Pan-KRAS inhibitor disables oncogenic signalling and tumour growth. Nature 619, 160–166. 10.1038/s41586-023-06123-3.

36. Salmon, M., Alvarez-Diaz, R., Fustero-Torre, C., Brehey, O., Lechuga, C.G., Sanclemente, M., Fernandez-Garcia, F., Lopez-Garcia, A., Martin-Guijarro, M.C., Rodriguez-Perales, S., et al. (2023). Kras oncogene ablation prevents resistance in advanced lung adenocarcinomas. J Clin Invest 133. 10.1172/JCI164413.

37. Hallin, J., Bowcut, V., Calinisan, A., Briere, D.M., Hargis, L., Engstrom, L.D., Laguer, J., Medwid, J., Vanderpool, D., Lifset, E., et al. (2022). Anti-tumor efficacy of a potent and selective non-covalent KRAS. Nat Med 28, 2171–2182. 10.1038/s41591-022-02007-7.

38. McDonald, D.M., and Choyke, P.L. (2003). Imaging of angiogenesis: from microscope to clinic. Nat Med 9, 713–725. 10.1038/nm0603-713.

39. Motta, M., Fidan, M., Bellacchio, E., Pantaleoni, F., Schneider-Heieck, K., Coppola, S., Borck, G., Salviati, L., Zenker, M., Cirstea, I.C., and Tartaglia, M. (2019). Dominant Noonan syndrome-causing LZTR1 mutations specifically affect the Kelch domain substrate-recognition surface and enhance RAS-MAPK signaling. Hum Mol Genet 28, 1007–1022. 10.1093/hmg/ddy412.

40. Pedley, R.B., El-Emir, E., Flynn, A.A., Boxer, G.M., Dearling, J., Raleigh, J.A., Hill, S.A., Stuart, S., Motha, R., and Begent, R.H. (2002). Synergy between vascular targeting agents and antibody-directed therapy. Int J Radiat Oncol Biol Phys 54, 1524–1531. 10.1016/s0360-3016(02)03923-8.

41. Baldelli, E., El Gazzah, E., Moran, J.C., Hodge, K.A., Manojlovic, Z., Bassiouni, R., Carpten, J.D., Ludovini, V., Baglivo, S., Crinò, L., et al. (2021). Wild-Type KRAS Allele Effects on Druggable Targets in KRAS Mutant Lung Adenocarcinomas. Genes (Basel) 12. 10.3390/genes12091402.

42. Sumita, K., Yoshino, H., Sasaki, M., Majd, N., Kahoud, E.R., Takahashi, H., Takeuchi, K., Kuroda, T., Lee, S., Charest, P.G., et al. (2014). Degradation of activated K-Ras orthologue via K-Ras-specific lysine residues is required for cytokinesis. J Biol Chem 289, 3950–3959. 10.1074/jbc.M113.531178.

43. Marzahn, M.R., Marada, S., Lee, J., Nourse, A., Kenrick, S., Zhao, H., Ben-Nissan, G., Kolaitis, R.M., Peters, J.L., Pounds, S., et al. (2016). Higher-order oligomerization promotes localization of SPOP to liquid nuclear speckles. EMBO J 35, 1254–1275. 10.15252/embj.201593169.

44. Li, C., Qin, T., Jin, Y., Hu, J., Yuan, F., Cao, Y., and Duan, C. (2023). Cerebrospinal fluid-derived extracellular vesicles after spinal cord injury promote vascular regeneration via PI3K/AKT signaling pathway. J Orthop Translat 39, 124–134. 10.1016/j.jot.2023.02.001.

45. Fleuren, E.D., Versleijen-Jonkers, Y.M., Roeffen, M.H., Franssen, G.M., Flucke, U.E., Houghton, P.J., Oyen, W.J., Boerman, O.C., and van der Graaf, W.T. (2014). Temsirolimus combined with cisplatin or bevacizumab is active in osteosarcoma models. Int J Cancer 135, 2770–2782. 10.1002/ijc.28933.

46. Damnernsawad, A., Bottomly, D., Kurtz, S.E., Eide, C.A., McWeeney, S.K., Tyner, J.W., and Nechiporuk, T. (2022). Genome-wide CRISPR screen identifies regulators of MAPK and MTOR pathways mediating sorafenib resistance in acute myeloid leukemia. Haematologica 107, 77–85. 10.3324/haematol.2020.257964.

47. Adachi, Y., Kimura, R., Hirade, K., Yanase, S., Nishioka, Y., Kasuga, N., Yamaguchi, R., and Ebi, H. (2023). Scribble mis-localization induces adaptive resistance to KRAS G12C inhibitors through feedback activation of MAPK signaling mediated by YAP-induced MRAS. Nat Cancer 4, 829–843. 10.1038/s43018-023-00575-2.

48. Wei, W., Geer, M.J., Guo, X., Dolgalev, I., Sanjana, N.E., and Neel, B.G. (2023). Genome-wide CRISPR/Cas9 screens reveal shared and cell-specific mechanisms of resistance to SHP2 inhibition. J Exp Med 220. 10.1084/jem.20221563.

49. Corcoran, R.B. (2023). A single inhibitor for all KRAS mutations. Nat Cancer 4, 1060–1062. 10.1038/s43018-023-00615-x.

50. Manning, B.D., and Toker, A. (2017). AKT/PKB Signaling: Navigating the Network. Cell 169, 381–405. 10.1016/j.cell.2017.04.001.

51. Kovalski, J.R., Bhaduri, A., Zehnder, A.M., Neela, P.H., Che, Y., Wozniak, G.G., and Khavari, P.A. (2019). The Functional Proximal Proteome of Oncogenic Ras Includes mTORC2. Mol Cell 73, 830–844.e812. 10.1016/j.molcel.2018.12.001.

## References

1. Steklov, M., Pandolfi, S., Baietti, M.F., Batiuk, A., Carai, P., Najm, P., Zhang, M., Jang, H., Renzi, F., Cai, Y., et al. (2018). Mutations in LZTR1 drive human disease by dysregulating RAS ubiquitination. Science 362, 1177–1182. 10.1126/science.aap7607.

2. Murgaski, A., Kiss, M., Van Damme, H., Kancheva, D., Vanmeerbeek, I., Keirsse, J., Hadadi, E., Brughmans, J., Arnouk, S.M., Hamouda, A.E.I., et al. (2022). Efficacy of CD40 Agonists Is Mediated by Distinct cDC Subsets and Subverted by Suppressive Macrophages. Cancer Res 82, 3785–3801. 10.1158/0008-5472.CAN-22-0094.

3. Marien, E., Hillen, A., Vanderhoydonc, F., Swinnen, J.V., and Vande Velde, G. (2017). Longitudinal microcomputed tomography-derived biomarkers for lung metastasis detection in a syngeneic mouse model: added value to bioluminescence imaging. Lab Invest 97, 24–33. 10.1038/labinvest.2016.114.

4. Berghen, N., Dekoster, K., Marien, E., Dabin, J., Hillen, A., Wouters, J., Deferme, J., Vosselman, T., Tiest, E., Lox, M., et al. (2019). Radiosafe micro-computed tomography for longitudinal evaluation of murine disease models. Sci Rep 9, 17598. 10.1038/s41598-019-53876-x.

5. Ivanisevic, T., Steklov, M., Lechat, B., Cawthorne, C., Gsell, W., Velde, G.V., Deroose, C., Van Laere, K., Himmelreich, U., Sewduth, R.N., and Sablina, A.A. (2023). Targeted STAT1 therapy for LZTR1-driven peripheral nerve sheath tumor. Cancer Commun (Lond) 43, 1386–1390. 10.1002/cac2.12490.

6. Descamps, B., Sewduth, R., Ferreira Tojais, N., Jaspard, B., Reynaud, A., Sohet, F., Lacolley, P., Allières, C., Lamazière, J.M., Moreau, C., et al. (2012). Frizzled 4 regulates arterial network organization through noncanonical Wnt/planar cell polarity signaling. Circ Res 110, 47–58. 10.1161/CIRCRESAHA.111.250936.

7. Sewduth, R.N., Kovacic, H., Jaspard-Vinassa, B., Jecko, V., Wavasseur, T., Fritsch, N., Pernot, M., Jeaningros, S., Roux, E., Dufourcq, P., et al. (2017). PDZRN3 destabilizes endothelial cell-cell junctions through a PKCζ-containing polarity complex to increase vascular permeability. Sci Signal 10. 10.1126/scisignal.aag3209.

8. McFall, T., Diedrich, J.K., Mengistu, M., Littlechild, S.L., Paskvan, K.V., Sisk-Hackworth, L., Moresco, J.J., Shaw, A.S., and Stites, E.C. (2019). A systems mechanism for KRAS mutant allele-specific responses to targeted therapy. Sci Signal 12. 10.1126/scisignal.aaw8288.

9. Stites, E.C., Trampont, P.C., Ma, Z., and Ravichandran, K.S. (2007). Network analysis of oncogenic Ras activation in cancer. Science 318, 463–467. 10.1126/science.1144642.

10. McFall, T., and Stites, E.C. (2021). Identification of RAS mutant biomarkers for EGFR inhibitor sensitivity using a systems biochemical approach. Cell Rep 37, 110096. 10.1016/j.celrep.2021.110096.

