## Supplementary figures and images for "Increased dosage of wild-type KRAS protein drives *KRAS*-mutant lung tumorigenesis and drug resistance"

### Supplementary Figures 1-3

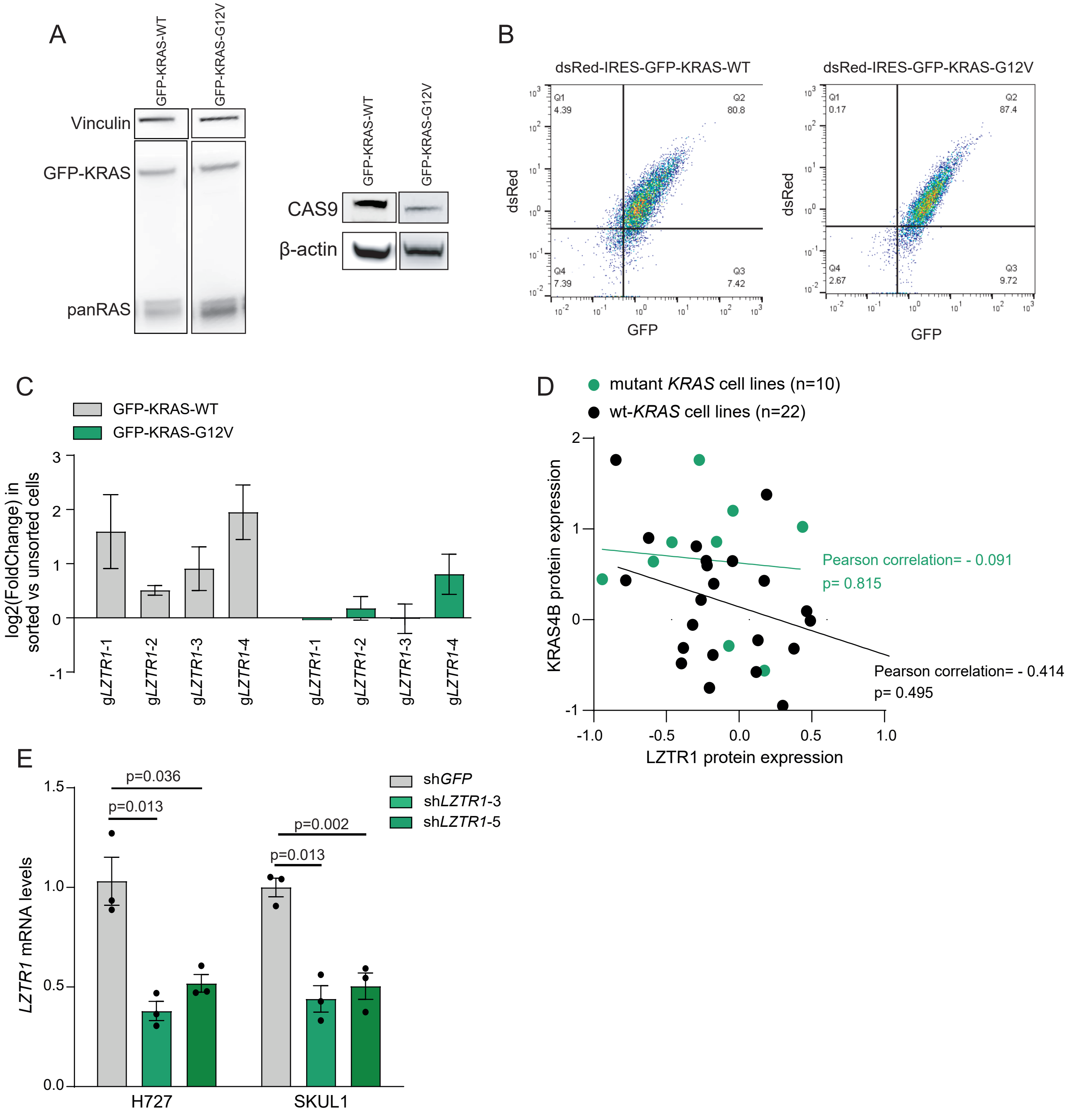

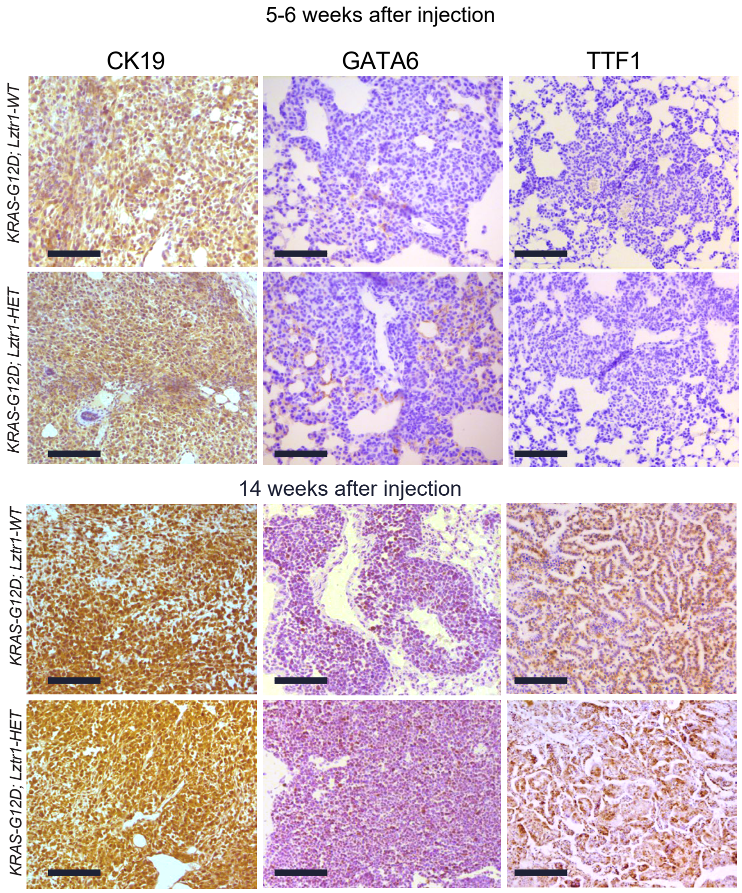

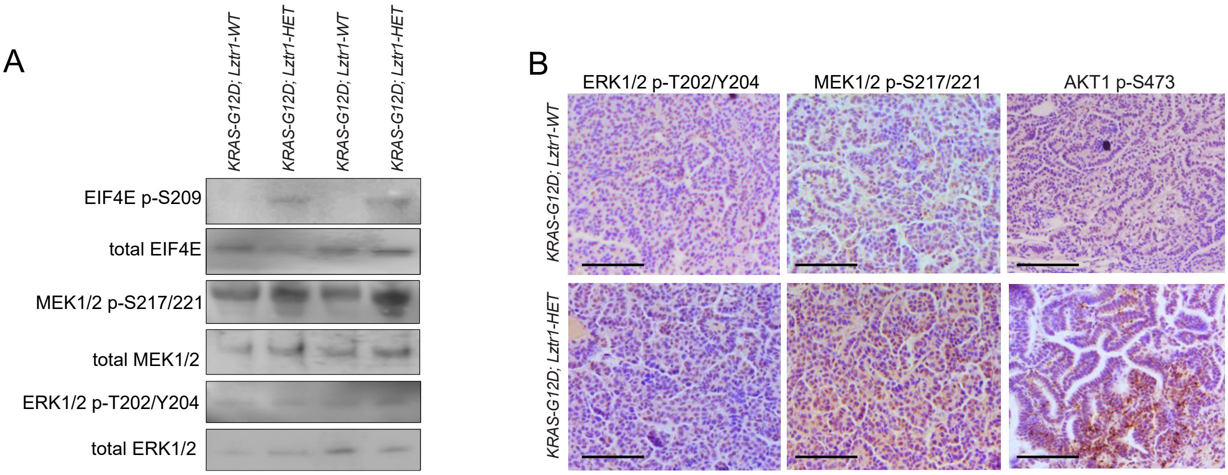
